# The ChvG–ChvI and NtrY–NtrX two-component systems coordinately regulate growth of *Caulobacter crescentus*

**DOI:** 10.1101/2021.04.17.440287

**Authors:** Benjamin J. Stein, Aretha Fiebig, Sean Crosson

## Abstract

Two-component signaling systems (TCSs) are comprised of a sensory histidine kinase and a response regulator protein. In response to environmental changes, sensor kinases directly phosphorylate their cognate response regulator to affect gene expression. Bacteria typically express multiple TCSs that are insulated from one another and regulate distinct physiological processes. There are certainly examples of cross-regulation between TCSs, but this phenomenon remains relatively unexplored. We have identified regulatory links between the ChvG–ChvI (ChvGI) and NtrY–NtrX (NtrYX) TCSs, which control important and often overlapping processes in α-proteobacteria, including maintenance of the cell envelope. Deletion of *chvG* and *chvI* in *Caulobacter crescentus* limited growth in defined medium and a selection for genetic suppressors of this growth phenotype uncovered interactions among *chvGI, ntrYX*, and *ntrZ*, which encodes a previously uncharacterized periplasmic protein. Significant overlap in the experimentally-defined ChvI and NtrX transcriptional regulons provided support for the observed genetic connections between *ntrYX* and *chvGI*. Moreover, we present evidence that the growth defect of strains lacking *chvGI* is influenced by the phosphorylation state of NtrX and, to some extent, by levels of the TonB-dependent receptor ChvT. Measurements of NtrX phosphorylation *in vivo* indicated that NtrZ is an upstream regulator of NtrY, and that NtrY primarily functions as an NtrX phosphatase. We propose a model in which NtrZ functions in the periplasm to inhibit NtrY phosphatase activity; regulation of phosphorylated NtrX levels by NtrZ and NtrY provides a mechanism to modulate and balance expression of the NtrX and ChvI regulons under different growth conditions.

**Importance:** Two-component signaling systems (TCSs) enable bacteria to regulate gene expression in response to physiochemical changes in their environment. The ChvGI and NtrYX TCSs regulate diverse pathways associated with pathogenesis, growth, and cell envelope function in many α-proteobacteria. We used *Caulobacter crescentus* as a model to investigate regulatory connections between ChvGI and NtrYX. Our work defined the ChvI transcriptional regulon in *C. crescentus* and revealed a genetic interaction between ChvGI and NtrYX, whereby modulation of NtrYX signaling affects the survival of cells lacking ChvGI. In addition, we identified NtrZ as a periplasmic inhibitor of NtrY phosphatase activity *in vivo*. Our work establishes *C. crescentus* as an excellent model to investigate multi-level regulatory connections between ChvGI and NtrYX in α-proteobacteria.

## Introduction

Bacteria employ two-component signaling systems (TCSs) to respond to environmental cues and maintain cellular homeostasis (1). TCS sensory modules consist of two core components: a sensory histidine kinase (HK) and a response regulator (RR). In response to signal(s), the HK undergoes autophosphorylation on a conserved histidine residue, and then passes the phosphoryl group to a conserved aspartate on the receiver (REC) domain of the RR (1). HKs may also act as phosphatases, dephosphorylating phosphorylated RRs (1). RR phosphorylation generally alters the activity of effector domains that change gene expression. TCSs are modular, and the output of a particular RR may vary between different organisms (2-5). TCSs that regulate host interactions in pathogens and symbionts are often conserved in related free-living organisms and enable responses to similar physiochemical cues present in the environment (2, 6-8).

Although typical TCSs rely on a single HK and RR pair, many systems incorporate additional proteins, such as activators or inhibitors, to form more complex signaling networks (9-17). Historically, most TCSs have been considered to be insular systems, but in many bacteria, cross-regulation between HKs and RRs may integrate multiple environmental cues (9, 11, 14, 18-20). Even when HK and RR pairs are well insulated, TCSs can interact at the transcriptional level (20, 21). For example, one TCS may regulate the expression of other TCS genes, or multiple TCSs may influence transcription of the same downstream gene (22-24).

ChvG–ChvI (ChvGI) and NtrY–NtrX (NtrYX) are conserved α-proteobacterial TCSs that often regulate similar physiological processes, raising the possibility that they may work together in a coordinated fashion (25, 26). The ChvG HK and ChvI RR were originally identified as pleiotropic regulators in the plant pathogen *Agrobacterium tumefaciens*, affecting virulence, detergent tolerance, and pH sensitivity (8, 27). Subsequent work has linked ChvGI to host interaction, cell motility, acid sensing, and exopolysaccharide production in a variety of α-proteobacteria (6, 7, 28-32). In most characterized systems, the periplasmic protein ExoR binds to ChvG and inhibits its kinase activity (15, 16, 33). Acidic pH activates the ChvGI system by triggering rapid proteolysis of ExoR (34). However, not all organisms with ChvGI, including *Caulobacter crescentus*, encode an ortholog of ExoR. These bacteria must regulate ChvG kinase activity by a different mechanism.

Like ChvGI, NtrYX (consisting of the NtrY HK and NtrX RR) is conserved in many α-proteobacteria, including multiple pathogens and symbionts (35-39). Although early studies concluded that NtrYX regulates nitrogen metabolism, recent work suggests that, in certain α-proteobacteria, it also controls exopolysaccharide biosynthesis, cell motility, and cell envelope composition (25, 35, 39, 40-44). Given that these processes are also regulated by ChvGI, it is conceivable that ChvGI and NtrYX act coordinately. However, no work to date has identified a substantial genetic interaction between these TCSs (25, 26).

*C. crescentus*, a free-living α-proteobacterium found in freshwater and soil environments, encodes both the ChvGI and NtrYX systems (45, 46). *C. crescentus* ChvGI activates transcription of the small regulatory RNA *chvR*, which post-transcriptionally represses the TonB-dependent receptor gene *chvT* (6). Examination of reporters of *chvR* transcription indicated that ChvGI is activated by growth in defined medium, acidic pH, DNA damage, growth at stationary phase, and cell envelope stress (6, 47). However, aside from *chvR*, genes regulated by ChvGI have not been defined. *C. crescentus* NtrYX is less well-characterized than ChvGI, but a recent study established that NtrX is phosphorylated in stationary phase in defined medium, as a result of acidification (48). In addition, NtrX appears to play a core role in regulating *C. crescentus* physiology, as *ntrX* is essential for growth in complex medium and *ΔntrX* cells grow more slowly than wild-type (WT) cells in defined medium (49).

In this study, we initially took a reverse genetic approach to characterize the role of ChvGI in regulating *C. crescentus* physiology. Deletion of *chvG* and *chvI* caused a distinctive growth defect in defined medium. By exploiting this defect, we identified striking genetic interactions between *chvGI* and *ntrY, ntrX*, and *ntrZ* (a previously uncharacterized gene). Epistasis analysis provided evidence that unphosphorylated NtrX is detrimental to cells lacking *chvG* or *chvI*. We defined the ChvI transcriptional regulon and discovered that it overlaps significantly with genes regulated by NtrX. In addition, we found that NtrZ promotes NtrX phosphorylation *in vivo*, likely by inhibiting NtrY phosphatase activity. We conclude that ChvGI and NtrYX interact at multiple transcriptional levels, working both in concert and in opposition to regulate *C. crescentus* growth in defined medium.

## Results

### Loss of the ChvGI system limits growth in defined medium

To investigate the physiological role of ChvGI in *C. crescentus*, we generated strains with in-frame deletions of *chvG* or *chvI* and examined their growth. The deletion strains grew normally in complex medium (peptone-yeast extract, PYE), but they displayed a distinctive growth defect in defined medium (M2 minimal salts medium with xylose as carbon source, M2X) (Fig. 1A, 1B and Fig. S1). Although overnight cultures of each strain inoculated from PYE agar plates grew to similar densities in M2X, strains lacking *chvG* or *chvI* exhibited reduced growth capacity after dilution, reaching a terminal density of only OD_660_ ∼ 0.1 (Fig. 1A). This lower cell density correlated with fewer colony-forming units (CFUs) (Fig. 1B). Ectopic overexpression of *chvG* (*chvG++)* or *chvI* (*chvI++)* from a xylose-inducible promoter (xylose is the sole carbon source in M2X medium) fully rescued growth of *ΔchvG* or *ΔchvI* strains, respectively, under these conditions (Fig. 1A, 1B).

**Figure 1:**
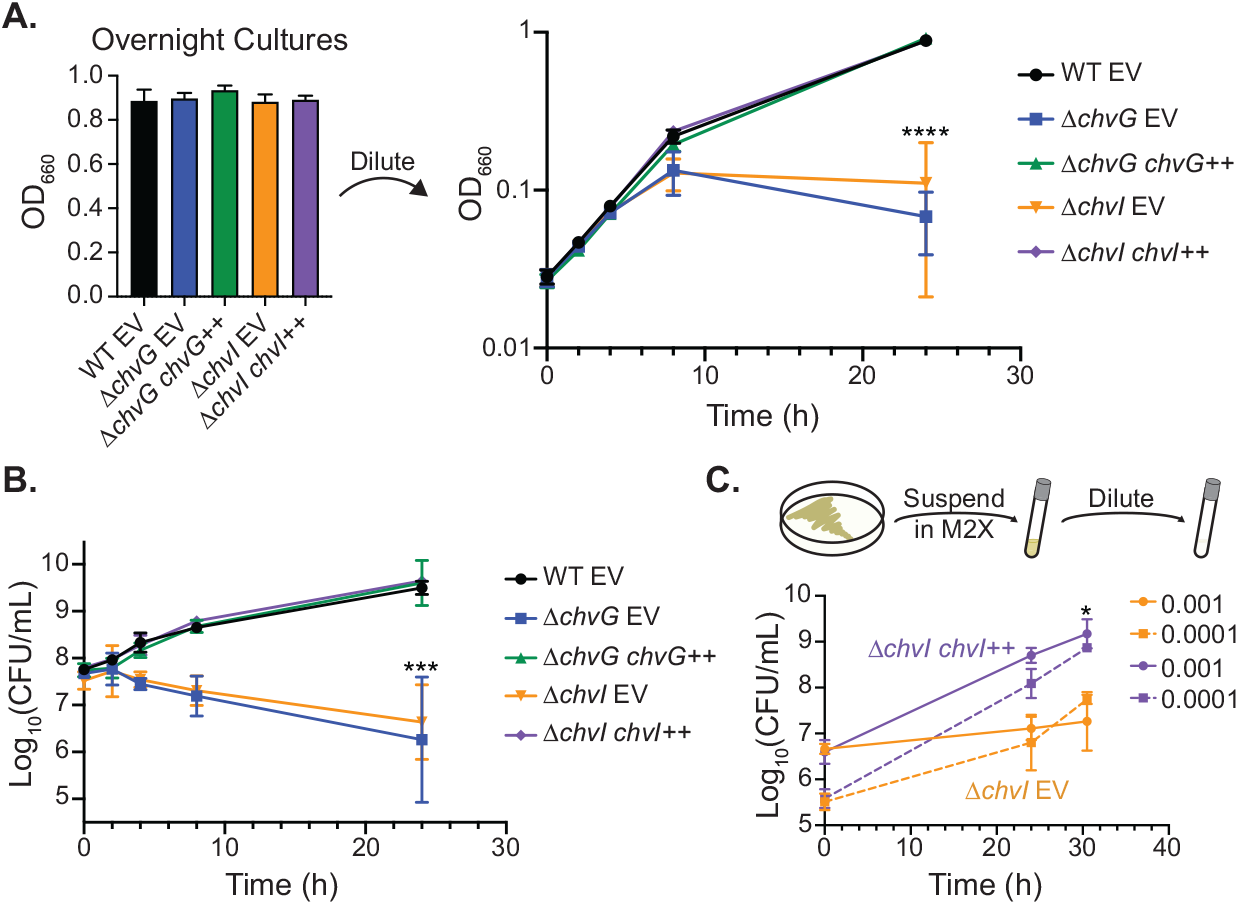
Loss of *chvG* or *chvI* limits culture density in defined M2X medium. (A) Growth curves, measured by optical density (OD_660_), of WT, *ΔchvG*, and *ΔchvI* strains bearing empty vector (EV) or genetic rescue plasmid (++) integrated at the xylose locus. Primary M2X cultures, inoculated from PYE plates, all grow to high density (left). However, upon back-dilution to OD_660_ = 0.025, *ΔchvG* and *ΔchvI* EV strains saturate at a significantly lower OD (right). Points represent averages of three biological replicates ± SD. ^****^ = *p* < 0.0001, one-way ANOVA followed by Dunnett’s post-test comparison to WT EV at 24 h. (B) Growth curves, measured by CFU, corresponding to the cultures in A. Points represent averages of three biological replicates SD. ^***^ = *p* < 0.0005, one-way ANOVA followed by Dunnett’s post-test comparison to WT EV at 24 h. (C) Growth of cultures inoculated from PYE agar plates at different starting densities, measured by CFU. *ΔchvI* cells carrying empty vector (EV) or genetic rescue plasmid (++) were suspended in M2X medium and immediately diluted to OD_660_ = 0.001 or 0.0001. Points represent averages of three biological replicates ± SD. ^*^ = *p* < 0.05, one-way ANOVA followed by Dunnett’s post-test comparison to *ΔchvI* EV, 10^−4^ dilution at 30.5h.

To evaluate if the growth defect was related to the number of cell divisions in defined medium, we resuspended cells in M2X from PYE agar plates and diluted cultures to several low starting densities. *ΔchvI* EV cultures started at OD_660_ = 0.001 and 0.0001 saturated at similar CFU values (∼10^7^ CFU/mL) that were significantly lower than those of *ΔchvI chvI++* cultures (∼10^9^ CFU/mL) (Fig. 1C). Thus, *ΔchvI* mutants that initiate from low density are not limited in the number of times they can divide in M2X medium, but rather reach a defined carrying capacity. *ΔchvI* cultures started at higher densities (OD_600_ = 0.05 and 0.1) grew to ∼10^9^ CFU/mL, indicating that a high starting density enables primary overnight cultures to saturate at higher density (Fig. S2A).

Washing *ΔchvI* cells once or twice with M2X before dilution had no effect on the number of CFUs at 30.5 h, suggesting that trace contaminating PYE components do not contribute to growth in M2X primary cultures (Fig. S2B). Moreover, cell density alone did not determine the ability of *ΔchvI* cultures to grow in M2X, as denser back-dilutions from *ΔchvI* overnight cultures did not reach higher viable cell counts (Fig. S2C). Together, our results suggest that the diminished growth capacity of *ΔchvI* or *ΔchvG* cells in M2X medium is a function of the time since resuspension from PYE plates and growth phase.

### ChvI phosphorylation is critical for growth in defined medium

Given the similarity between the phenotypes displayed by *ΔchvI* and *ΔchvG* strains, we predicted that phosphorylation of ChvI might be important for growth in M2X medium. To test this hypothesis, we generated strains harboring *chvI* alleles encoding changes to the conserved sites of phosphorylation (D52A, D52N, and D52E). Growth of strains carrying the non-phosphorylatable alleles *chvI(D52A)* and *chvI(D52N)* was similar to *ΔchvI* cells in M2X (Fig. 2A and Fig. S3A). By contrast, *chvI(D52E)* cultures reached higher densities than *ΔchvI* cultures, suggesting that ChvI(D52E) is active, albeit not to the same level as phosphorylated ChvI (Fig. 2A and Fig. S3A). Substitutions of the phosphorylatable aspartate with glutamate often act as phosphomimetic mutations and constitutively activate RRs (50-53). Thus, this result supports a critical role for ChvI phosphorylation during growth in M2X medium.

**Figure 2:**
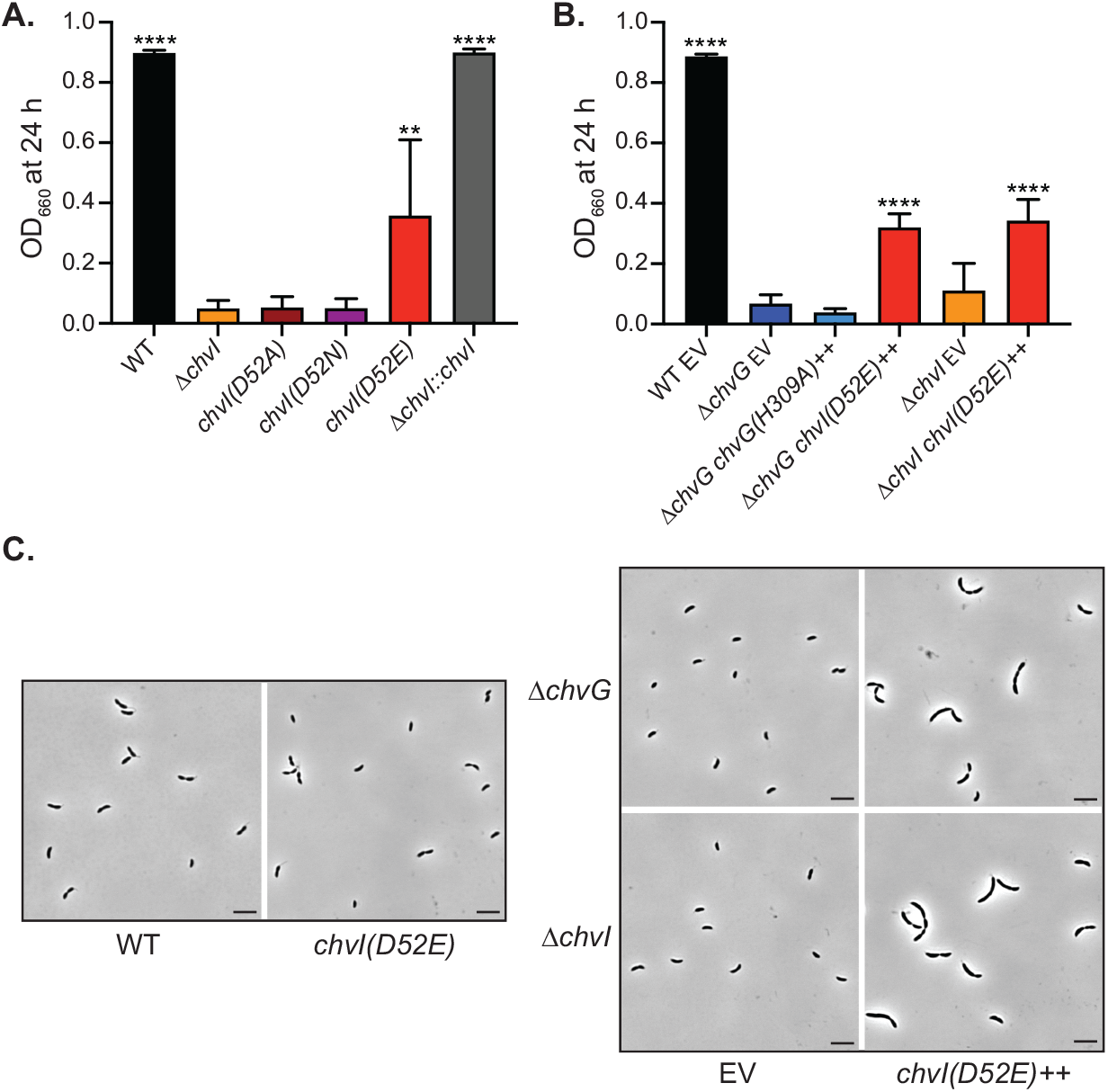
The *chvI(D52E)* allele is active and induces filamentation in M2X medium when overexpressed. (A) Optical density (OD_660_) of cultures 24 h after back-dilution to OD_660_= 0.025 in M2X medium (Average ± SD, *N* 4). *chvI* mutant strains were generated by restoring the *ΔchvI* allele with point-mutant alleles through allelic exchange. Similarly, restored WT (*ΔchvI*::*chvI*) was generated by replacement of the *ΔchvI* locus with WT *chvI*. ^*^ = *p* < 0.05, ^**^ = *p* < 0.01, ^****^ *p* < 0.0001 one-way ANOVA followed by Dunnett’s post-test comparison to *ΔchvI*. (B) Optical density (OD_660_) of strains carrying empty vector (EV) or overexpression plasmids (++) 24 h after back-dilution to OD_660_ = 0.025 in M2X medium (average ± SD, *N* = 3). ^****^ = *p* < 0.0001, one-way ANOVA followed by Dunnett’s post-test comparison to *ΔchvG* EV. (C) Phase-contrast micrographs of primary overnight cultures grown in M2X medium. On the right, *ΔchvI* and *ΔchvG* strains carry empty vector (EV) or overexpression plasmids (++). Scale bars, 5 µm.

To further examine the importance of ChvI phosphorylation, we overexpressed *chvG* and *chvI* alleles from a xylose-inducible promoter in knockout backgrounds. Although overexpression of *chvG* rescued growth of *ΔchvG* cells (Fig 1B), overexpression of *chvG(H309A)*, a catalytic-histidine mutant, did not, indicating that phosphorylation of ChvG, and by extension ChvI, is important for growth in M2X medium (Fig. 2B). Overexpression of *chvI(D52E)* only partially rescued growth of *ΔchvI* and *ΔchvG* strains in M2X and these strains grew similarly to the strain encoding *chvI(D52E)* at the native locus (Fig. 2A, 2B and Fig. S3B). However, in contrast to native expression of *chvI(D52E)*, overexpression of this allele resulted in cell filamentation in M2X, but not PYE, consistent with a cell division defect (Fig. 2C and Fig. S3C). Thus, overexpression of phosphomimetic *chvI(D52E)* appears to interfere with cell division and, perhaps as a result, only partially rescues growth in M2X medium.

### Mutations in *ntrY, ntrX*, and a gene of unknown function, rescue the growth defect of *ΔchvI* and *ΔchvG* strains

To better understand why ChvGI is critical for growth in defined medium, we employed a selection strategy to isolate second-site mutations that alleviate the growth defect of *ΔchvG* and *ΔchvI* cells.

*ΔchvG* and *ΔchvI* M2X*-*overnight cultures were back-diluted and grown until they reached high density, presumably due to proliferation of suppressor strains (Fig. 3A). We then isolated single colonies with WT-like growth in M2X and used whole-genome sequencing to identify any acquired mutations (Fig. 3B).

**Figure 3:**
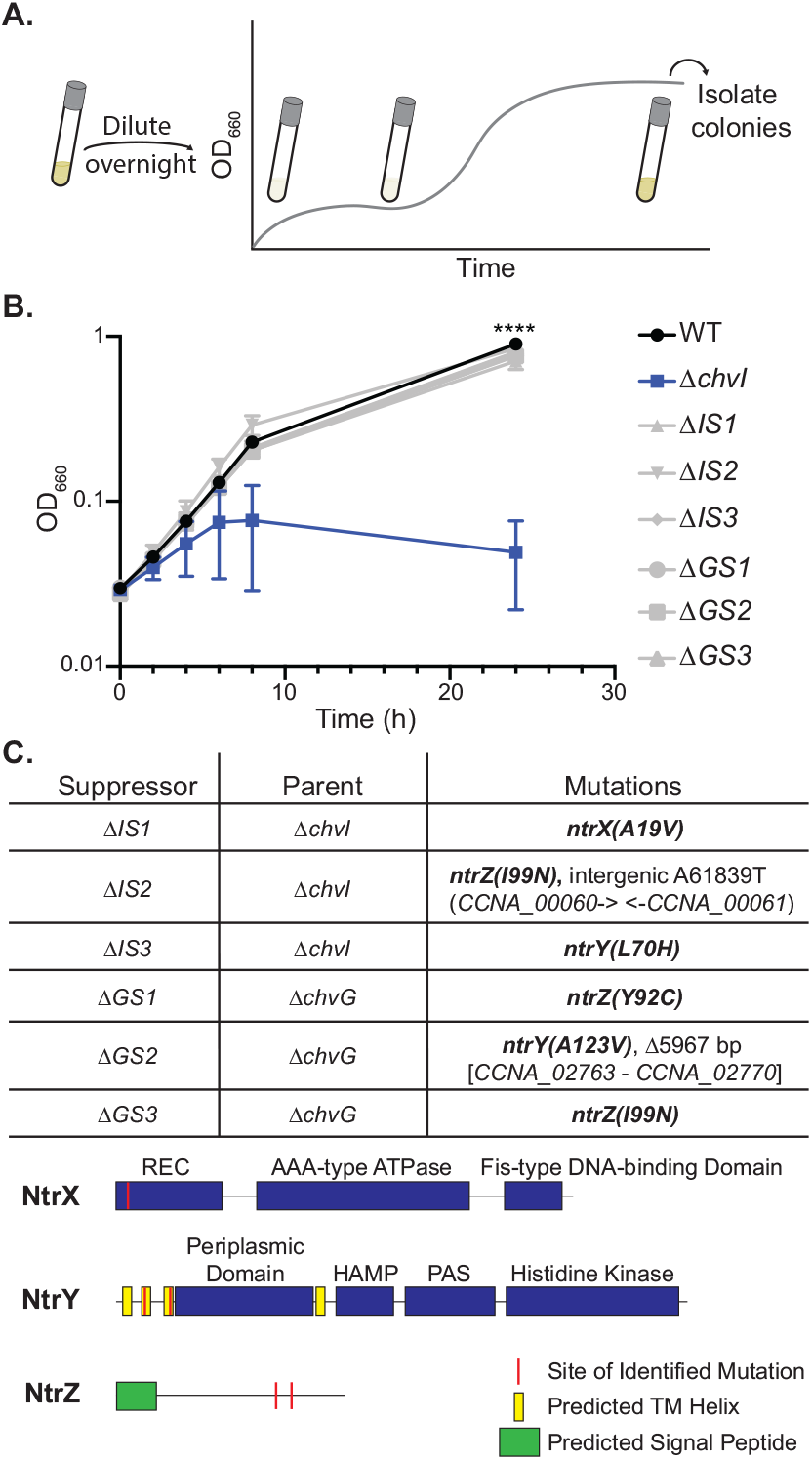
Mutations in the NtrYX TCS and a gene of unknown function suppress the growth defect of *ΔchvG* and *ΔchvI* strains in M2X medium. (A) Schematic representation of the suppressor selection protocol. Primary overnight cultures in M2X medium were back-diluted to OD_660_ = 0.025 and grown until cultures grew to high turbidity. Single colonies were isolated for confirmation and sequencing. (B) Growth curves, measured by optical density (OD_660_), of WT, *ΔchvI*, and suppressor strains isolated from either the *ΔchvI* strain (*ΔIS#*) or the *ΔchvG* strain (*ΔGS#*) upon back-dilution in M2X medium. Points represent averages of three biological replicates SD. ^****^ = *p* < 0.0001, one-way ANOVA followed by Dunnett’s post-test comparison to *ΔchvI* at 24 h. (C) Whole-genome sequencing results of each suppressor strain. The suppressor strain, parental strain, and identified polymorphism(s) are indicated. Mutations in *ntrY, ntrX*, and *ntrZ* (*CCNA_03863*) are in bold. Domain structures of NtrY, NtrX, and NtrZ are diagramed with domains in blue, signal peptides in green, transmembrane helices in yellow, and identified mutations in red.

In multiple independent strains, we identified nonsynonymous substitutions in the genes encoding the HK NtrY and its cognate RR NtrX (Fig. 3C). The *ntrY* mutations (L70H and A123V) are located in predicted transmembrane helices, which are involved in transmitting information from periplasmic sensing domains to the kinase domain (Fig. 3C) (54-56). We identified only one *ntrX* allele, A19V, which lies in helix 1 (α1) of the receiver domain (Fig. 3C). α1 is involved in the interaction interface between the HK dimerization and histidine phosphotransfer (DHp) domain and the RR REC domain (57, 58). We also note that the *ntrX(A19V)* strain was isolated on PYE plates and appeared to grow normally in M2X, suggesting that this is not a complete loss-of-function allele (49).

In addition to mutations in *ntrY* and *ntrX*, we identified three mutants with nonsynonymous substitutions in a gene encoding a putative periplasmic protein of unknown function (*CCNA_03863*, hereafter referred to as *ntrZ*) (Fig. 3C). The mutations identified (Y92C and I99N) are both located outside of the predicted signal sequence (Fig. 3C). Therefore, mutations in the NtrYX TCS and NtrZ appear to affect the growth of *ΔchvI* and *ΔchvG* cells in M2X medium.

### Overexpression or deletion of *ntrX* modulates the *ΔchvI* growth phenotype

Several studies have noted similar phenotypes of *chvGI* and *ntrYX* mutants (25, 26, 41); however, to our knowledge none have established a genetic link between these TCSs. To better understand the connections between ChvGI and NtrYX in *C. crescentus*, we first characterized the genetic relationship between *chvI* and *ntrX*. Replacement of *ntrX* with the *ntrX(A19V)* allele restored growth of *ΔchvI* cells in M2X (Fig. 4A), ruling out other background mutations as causal for suppression of the *ΔchvI* phenotype. However, overexpression of *ntrX(A19V)* in the presence of the native *ntrX* allele did not rescue growth of *ΔchvI* cells (Fig. 4A). This result indicates that the *ntrX(A19V)* allele is recessive, and suggests that the presence of WT NtrX contributes to the growth phenotype of *ΔchvI* cells.

**Figure 4:**
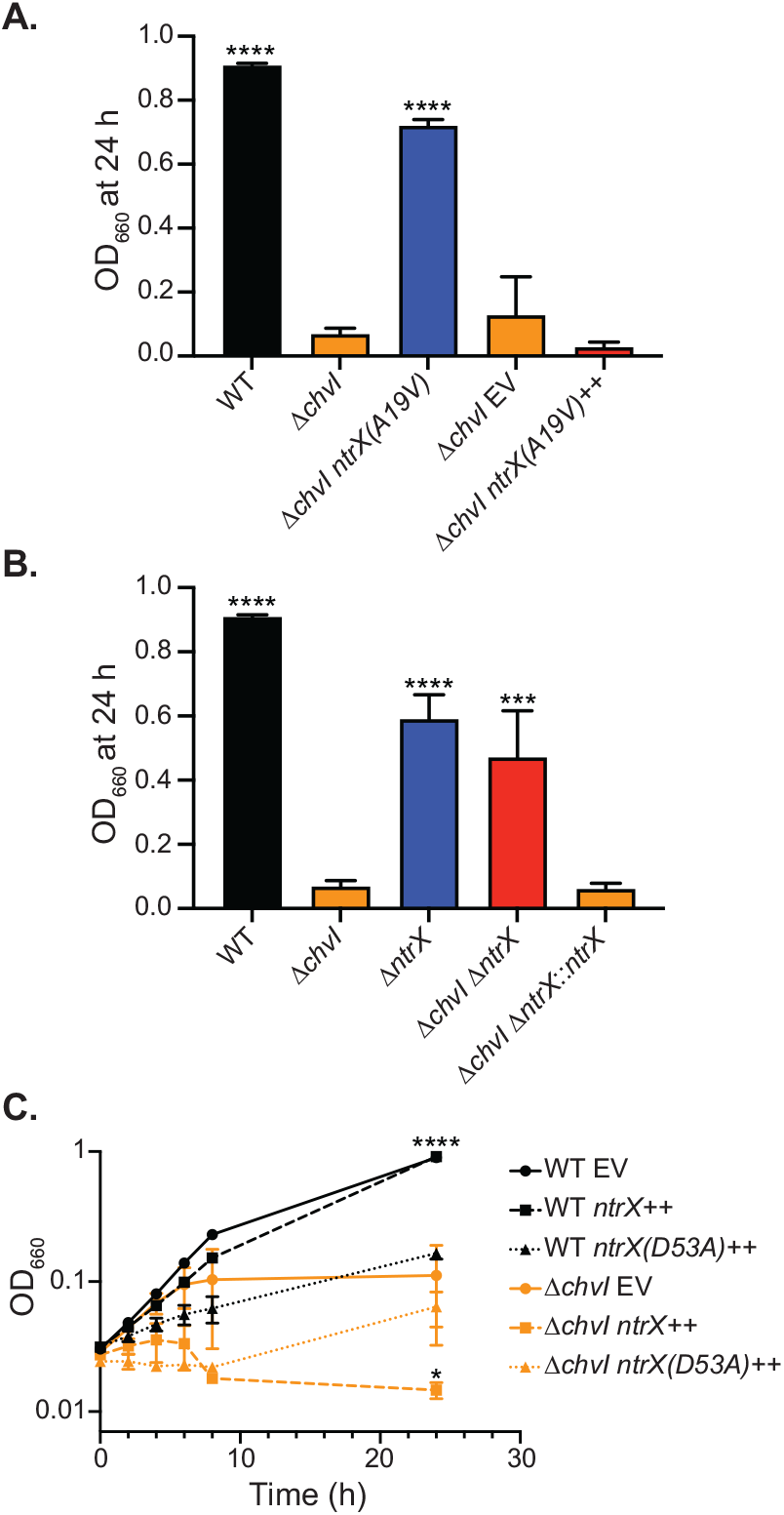
Non-phosphorylatable NtrX limits growth of cells in M2X medium. (A) Optical density (OD_660_) of cultures 24 h after back-dilution to OD_660_ = 0.025 in M2X medium (average ± SD, *N* = 3). The *ntrX(A19V)* allele was introduced by allele replacement for comparison with WT and *ΔchvI* or by overexpression (++) for comparison with an empty vector (EV) strain. ^****^ = *p* < 0.0001, one-way ANOVA followed by Dunnett’s post-test comparison to *ΔchvI*. (B) Optical density (OD_660_) of cultures 24 h after back-dilution to OD_660_ = 0.025 in M2X medium (average ± SD, *N* = 3). Restored *ΔchvI* (*ΔchvI ΔntrX*::*ntrX*) was generated by knocking *ntrX* into the native *ntrX* locus in *ΔchvI ΔntrX* cells. ^***^ = *p* < 0.0005, ^****^ = *p* < 0.0001, one-way ANOVA followed by Dunnett’s post-test comparison to *ΔchvI*. (C) Growth curves, measured by optical density (OD_660_), of strains upon back-dilution in M2X medium. WT and *ΔchvI* strains bear empty vector (EV) or *ntrX* overexpression vectors (++). Points represent averages of three biological replicates ± SD. ^*^ = *p* < 0.05, ^****^ = *p* < 0.0001, one-way ANOVA followed by Dunnett’s post-test comparison to *ΔchvI* EV at 24h.

To examine this notion further, we tested the effect of deleting *ntrX* in the *ΔchvI* strain. *ΔntrX* cells had a slight growth defect in M2X, but were clearly distinct from *ΔchvI* cells (Fig. 4B). Deletion of *ntrX* in a *ΔchvI* background rescued growth, indicating that the presence of *ntrX* is indeed detrimental to *ΔchvI* cells (Fig. 4B). Suppression of the *ΔchvI* growth phenotype was not due to a second-s ite mutation, as restoration of the *ntrX* locus (*ΔchvI ΔntrX*::*ntrX)* restored the growth defect in M2X medium (Fig. 4B).

Given that the presence of *ntrX* limits growth of *ΔchvI* cells in M2X, we next tested whether overexpression of *ntrX* affects growth of WT cells. Overexpression of *ntrX* only moderately slowed growth of WT cells. However, overexpression of the non-phosphorylatable *ntrX(D53A)* allele dramatically impaired the growth of WT cells in M2X, similar to the defect observed in *ΔchvI* cells (Fig. 4C). Overexpression of either *ntrX* or *ntrX(D53A)* exacerbated the growth defect of *ΔchvI* cells (Fig. 4C). Together these results support a model in which unphosphorylated NtrX limits growth capacity in M2X medium, especially in *ΔchvI* cells.

### NtrZ is a predicted periplasmic protein that functions upstream of NtrY to regulate levels of phosphorylated NtrX

We next examined the nature of the *ntrY* and *ntrZ* suppressor mutations. Overexpression of the *ntrY* and *ntrZ* alleles identified in our suppressor strains restored growth of *ΔchvI* cells in M2X (Fig. 5A). In addition, overexpression of WT *ntrY*, but not WT *ntrZ*, also significantly suppressed the *ΔchvI* growth defect (Fig. 5A). We conclude that the *ntrY* and *ntrZ* mutations are dominant, likely gain-of-function, alleles. As deletion of *ntrX* rescued the growth of *ΔchvI* cells, overexpression of *ntrY* or *ntrZ* mutant alleles likely promotes phosphorylation and/or sequestration of NtrX. We attempted, but failed, to construct *ΔchvI ΔntrY* and *ΔchvI ΔntrZ* double-mutants trains, suggesting that deletion of either *ntrY* or *ntrZ* is synthetically lethal with *chvI* deletion. This inability to isolate either double mutant also hinted that NtrY and NtrZ function in the same pathway. To test this possibility, we evaluated the phenotype of *ntrY* and *ntrZ* deletions in M2X medium. However, the growth of both single deletions and the Δ*ntrY ΔntrZ* double deletion was indistinguishable from WT, thus preventing epistasis analysis (Fig. 5B).

**Figure 5:**
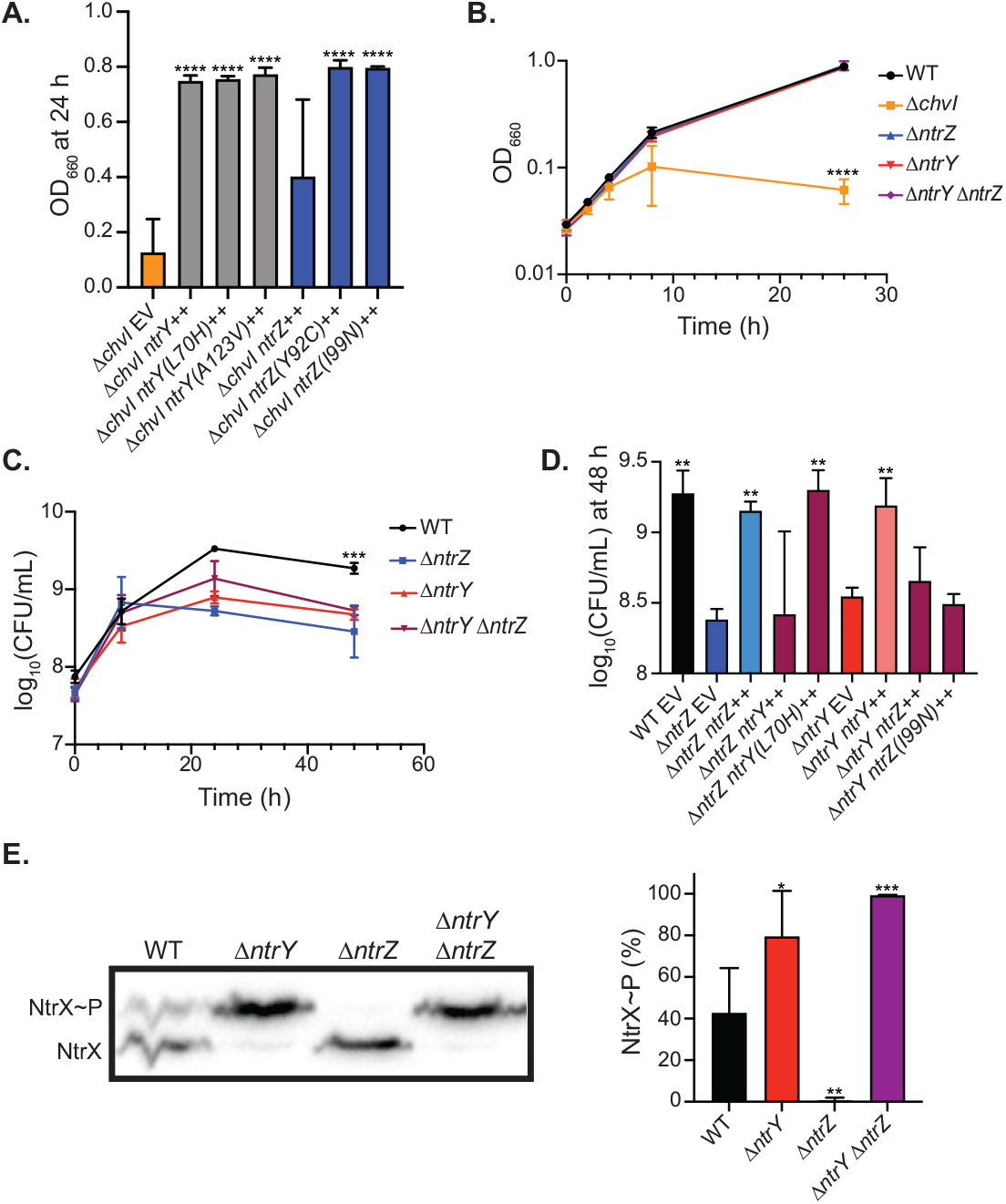
NtrZ functions upstream of NtrY as a phosphatase inhibitor. (A) Optical density (OD_660_) of strains bearing empty vector (EV) or *ntrZ* and *ntrY* overexpression vectors (++) 24 h after back-dilution to OD_660_ = 0.025 in M2X medium (average ± SD, *N* = 3). ^****^ = *p* < 0.0001, one-way ANOVA followed by Dunnett’s post-test comparison to *ΔchvI* EV. (B) Growth curves, measured by optical density (OD_660_), of WT and knockout strains upon back-dilution in M2X medium. Points are averages of three biological replicates ± SD. ^****^ = *p* < 0.0001, one-way ANOVA followed by Dunnett’s post-test comparison to WT at 24 h. (C) Growth curves, measured by CFU, for WT and mutant strains grown in M2X medium. Points are averages of three biological replicates ± SD. ^***^ = *p* = 0.0005, one-way ANOVA followed by Dunnett’s post-test comparison to *ΔntrZ* at 48 h. (D) Cell density, measured by CFU, for *ΔntrZ* and *ΔntrY* strains bearing empty vector (EV) or overexpression vectors (++) at 48 h growth in M2X medium. Points are averages of three biological replicates ± SD. ^**^ = *p* < 0.01, one-way ANOVA followed by Dunnett’s post-test comparison to *ΔntrZ* EV at 48 h. (E) Anti-HA western blot of lysates analyzed by Phos-tag gel electrophoresis (left). Each strain encodes *ntrX-HA* at the native *ntrX* locus. Phosphorylated (NtrX∼P) and unphosphorylated (NtrX) NtrX-HA are indicated. Quantification of the percent of NtrX that is phosphorylated (% NtrX∼P) (Right). Points are averages of four biological replicates ± SD. ^*^ = *p* < 0.05, ^**^ = *p* < 0.01, ^***^ = *p* < 0.001, one-way ANOVA followed by Dunnett’s post-test comparison to WT.

Phosphorylation of NtrX is likely important for stationary phase survival (48), and thus, we hypothesized that deletion of the *ntrY* HK, and perhaps also *ntrZ*, might lead to a stationary-phase defect. Indeed, we observed significantly lower CFUs in both *ΔntrY* and *ΔntrZ* cultures compared to WT cultures after 48 h of growth in M2X (Fig 5C). Notably, *ΔntrY ΔntrZ* cultures behaved similarly to the single mutants, suggesting that NtrY and NtrZ indeed function in the same pathway.

To place *ntrY* and *ntrZ* relative to one another, we evaluated the ability of *ntrY* and *ntrZ* alleles to rescue stationary-phase survival in the deletion strains. Both *ΔntrY* and Δ*ntrZ* cells were fully rescued by ectopic overexpression of each respective WT allele (Fig. 5D). Overexpression of *ntrY(L70H)*, but not *ntrY*, also fully rescued the phenotype of *ΔntrZ* cells (Fig. 5D). By contrast, neither WT *ntrZ* nor *ntrZ(I99N)* rescued *ΔntrY* cells (Fig. 5D). These data provide evidence that NtrZ functions upstream of NtrY, potentially as a kinase activator and/or phosphatase inhibitor.

We next examined the phosphorylation state of NtrX directly using Phos-tag gel electrophoresis. To detect NtrX by western blotting, we constructed strains carrying C-terminally HA-tagged *ntrX* (*ntrX*-*HA*) encoded at the native *ntrX* locus. Phos-tag analysis of WT lysates at stationary phase revealed distinct bands for phosphorylated (NtrX∼P) and unphosphorylated (NtrX) NtrX-HA (Fig. 5E). Deletion of *ntrZ* ablated NtrX∼P, consistent with a role for NtrZ in promoting NtrX phosphorylation (Fig. 5E). Surprisingly, *ΔntrY* cells displayed higher levels of NtrX phosphorylation than WT cells, suggesting that NtrY primarily acts as a phosphatase *in vivo* and is not required for NtrX phosphorylation. NtrZ indeed acts upstream of NtrY, as deletion of *ntrZ* in the *ΔntrY* background did not affect NtrX phosphorylation (Fig. 5E). We conclude that NtrZ promotes NtrX phosphorylation by inhibiting NtrY phosphatase activity.

Our characterization of the *ntrY, ntrX*, and *ntrZ* mutants suggested that phosphorylation of NtrX may rescue the growth of *ΔchvI* or *ΔchvG* cells. NtrX is phosphorylated under acidic conditions, such as those encountered during stationary phase in M2G or M2X medium (48). Therefore, we tested whether the pH of M2X affected the growth of *ΔchvI* cultures. For both WT and *ΔchvI* strains, primary overnight cultures diluted in M2X at pH 7.0 reached similar CFUs at 8 h as those diluted in standard M2X (pH 7.2) (Fig. 1B, S4). WT cultures were relatively unaffected by growth in M2X at pH 6.0, but had significantly fewer CFUs in M2X pH 5.5 vs. pH 7.0 (Fig. S4C). By contrast, *ΔchvI* cells were markedly less fit in M2X pH 6.0 than at pH 7.0, and displayed an intermediate growth yield at pH 5.5 relative to M2X at either pH 6.0 or pH 7.0 (Fig. S4C). These results are consistent both with suppression of the *ΔchvI* growth defect by NtrX phosphorylation and an important role for ChvGI in acid-stress responses (6).

### ChvI and NtrX regulate transcription of an overlapping set of genes

Although ChvGI is known to regulate the expression of *chvR*, the complete ChvI transcriptional regulon is not known. To examine the regulatory link between *ntrX* and *chvI* in greater detail, we performed an RNA-seq experiment to comprehensively define ChvI-dependent gene regulation. As *ΔchvI* cells grow poorly in M2X medium, we exploited overexpression of the phosphomimetic *chvI(D52E)* allele to assess ChvI-dependent transcription in PYE medium. Excluding the internal *chvI* control, we identified 162 genes with > 1.5 fold-change in *ΔchvI chvI(D52E)++* cells compared to *ΔchvI* EV cells (Fig. 6A and Table S1). Of those, 140 were upregulated and 22 were downregulated, indicating that ChvI primarily serves as a transcriptional activator. Consistent with previous work, *chvR* was upregulated while *chvT* was downregulated by overexpression of *chvI(D52E)* (6, 47). In addition, expression of both *chvG* and *hprK* was enhanced in cells expressing *chvI(D52E)*, pointing to positive autoregulation of the *chvIG-hprK* operon. We also observed regulation of multiple genes involved in envelope maintenance, metabolism, protein quality control, and transport (Fig. S5A). ChvI-dependent genes included multiple genes encoding proteases/peptidases (*CCNA_01341, CCNA_02846, CCNA_01955, CCNA_02721, mmpA, CCNA_02594, CCNA_01121, CCNA_01202*), peptidyl-prolyl and disulfide isomerases (*CCNA_02889, CCNA_01654, CCNA_01759, CCNA_01653, CCNA_00378, CCNA_00379*), members of the β-barrel assembly machine (BAM) complex (*bamA, bamB, bamD, bamE, bamF*), members of the Tol–Pal complex (*tolB, ybgF, tolQ*), and lipopolysaccharide biosynthesis genes (*CCNA_01497, CCNA_03454, CCNA_01496, lpxC*). Moreover, nearly 40% of ChvI regulon genes encoded hypothetical proteins or proteins of unknown function (Fig. S5A). MEME analysis of the top 35 upregulated operons identified a putative ChvI binding motif, with GCC direct repeats 11 nucleotides apart, that closely resembles the recently characterized binding motif of *Sinorhizobium meliloti* ChvI (Fig. S5B) (5).

**Figure 6:**
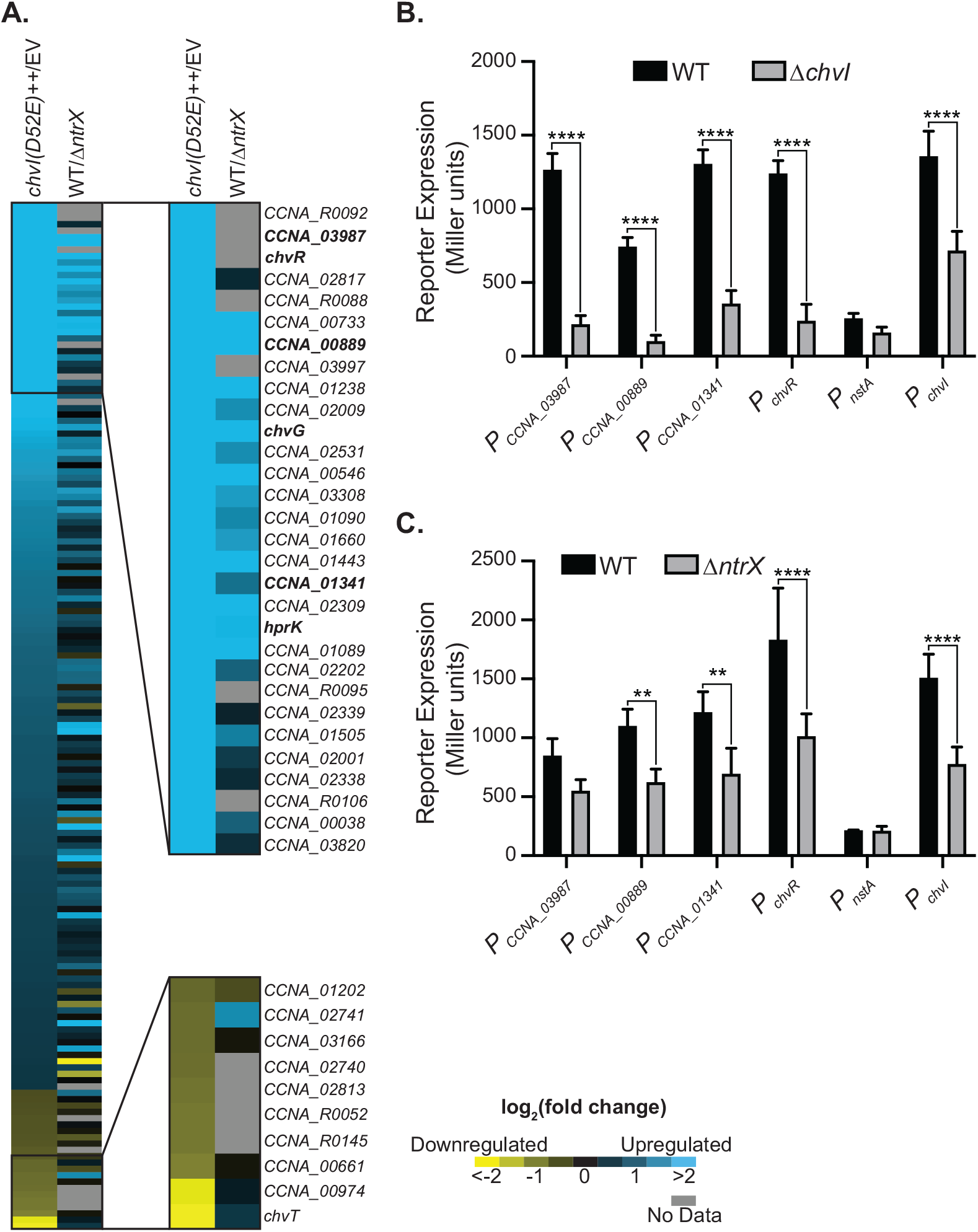
The ChvI and NtrX regulons overlap significantly. (A) Heat map of log_2_(fold change) for genes in the ChvI regulon (fold Change > 1.5, FDR *p-*value < 0.05) defined by RNA-seq (*ΔchvI chvI(D52E)*++ vs. *ΔchvI* EV). Log_2_(fold change) is also shown for a microarray dataset comparing RNA levels in WT and *ΔntrX* cells (49); grey cells indicate no data. The expression of upregulated genes (blue) is activated by ChvI or NtrX, whereas the expression of downregulated genes (yellow) is repressed by ChvI or NtrX. The genes most strongly regulated in the ChvI regulon are annotated (expression of those in bold is investigated further in B and C). (B) *lacZ* transcriptional reporter activity for a subset of genes in the ChvI regulon in WT and *ΔchvI* strains in M2X medium (average ± SD, *N* = 3). ^****^ = *p* < 0.0001, one-way ANOVA followed by Šídák’s post-test comparison for indicated pairs. (C) *lacZ* transcriptional reporter activity for the same genes as in B in WT and *ΔntrX* strains in M2X medium (average ± SD, *N* = 4). ^**^ = *p* < 0.01, ^****^ = *p* < 0.0001, one-way ANOVA followed by Šídák’s post-test comparison for indicated pairs.

As our regulon was defined by overexpression of a phosphomimetic ChvI mutant, we sought to validate our dataset under more physiological conditions. We constructed transcriptional β-galactosidase reporters for six selected regulon genes, from *nstA* (2-fold activation) to *CCNA_03987* (128-fold activation). Strains carrying these transcriptional reporters were initially grown in PYE medium, followed by back-dilution and 4 h of growth in M2X medium. Consistent with previous reports, the reporter for the *chvR* promoter (*P*_*chvR*_) was expressed in M2X in a *chvI*-dependent manner (Fig. 6B) (6, 47). The remaining reporters, apart from *P*_nstA_, which was expressed at low levels, also exhibited clear *chvI* dependence, supporting our RNA-seq results (Fig. 6B). Unlike the other reporters, *P*_*chvI*_ was only 2-fold lower in *ΔchvI* cells compared to WT cells, indicating that additional factors promote expression of the *chvIG-hprK* operon.

We next compared our ChvI regulon with a previously published NtrX regulon that was determined using DNA microarrays (49). This experiment measured relative gene expression between *C. crescentus* WT (strain NA1000) and *ΔntrX* cells during exponential growth in M2G medium, a condition where NtrX is expected to be largely unphosphorylated (48, 49). Surprisingly, a large fraction of the genes regulated by ChvI were also represented in the genes regulated by NtrX (Fig. 6A). That is, many of the genes upregulated by ChvI also appeared to be upregulated by NtrX (and likewise for downregulated genes). Using cutoffs of 1.5 fold-change for the ChvI RNA-seq data and 2.5 fold-change for the NtrX microarray data, we established that the ChvI regulon is significantly enriched for NtrX-dependent genes (6.31-fold enrichment, *p*-value = 8.99 × 10^−19^, hypergeometric test). To confirm this overlap, we evaluated the effect of deleting *ntrX* on our ChvI regulon reporters. Strains carrying the transcriptional reporters were grown to log phase (OD_660_ ∼ 0.1–0.2) in M2X medium before assaying β-galactosidase activity. Four of the six reporters exhibited *ntrX* dependence, including two genes (*CCNA_00889* and *chvR*) not evaluated in the NtrX microarray experiment (Fig. 6C). We note that transcription from the *chvI* promoter (*P*_*chvI*_) was activated by both *ntrX* and *chvI*, raising the possibility that NtrX may indirectly affect ChvI-dependent genes via upregulation of *chvIG-hprK*.

Given the oppositional nature of ChvI and NtrX during growth in M2X medium, we were surprised to see such a high degree of similarity in the genes they regulate. However, a small subset of genes exhibited opposite regulation by ChvI and NtrX, which might therefore account for suppression of the *ΔchvI* growth defect by deletion of *ntrX* (Fig. S6A). We overexpressed (for those genes upregulated by ChvI) or knocked out (for those genes downregulated by ChvI) each of these genes in *ΔchvI* cells and tested their growth capacity in M2X medium. Only one candidate, *chvT*, had any effect on *ΔchvI* cells. Specifically, deletion of *chvT* partially rescued growth of *ΔchvI* cells in M2X (Fig. 7A and Fig. S6B). Differential regulation of *chvT* expression by ChvI and NtrX may therefore contribute to the growth defect observed in M2X medium.

**Figure 7:**
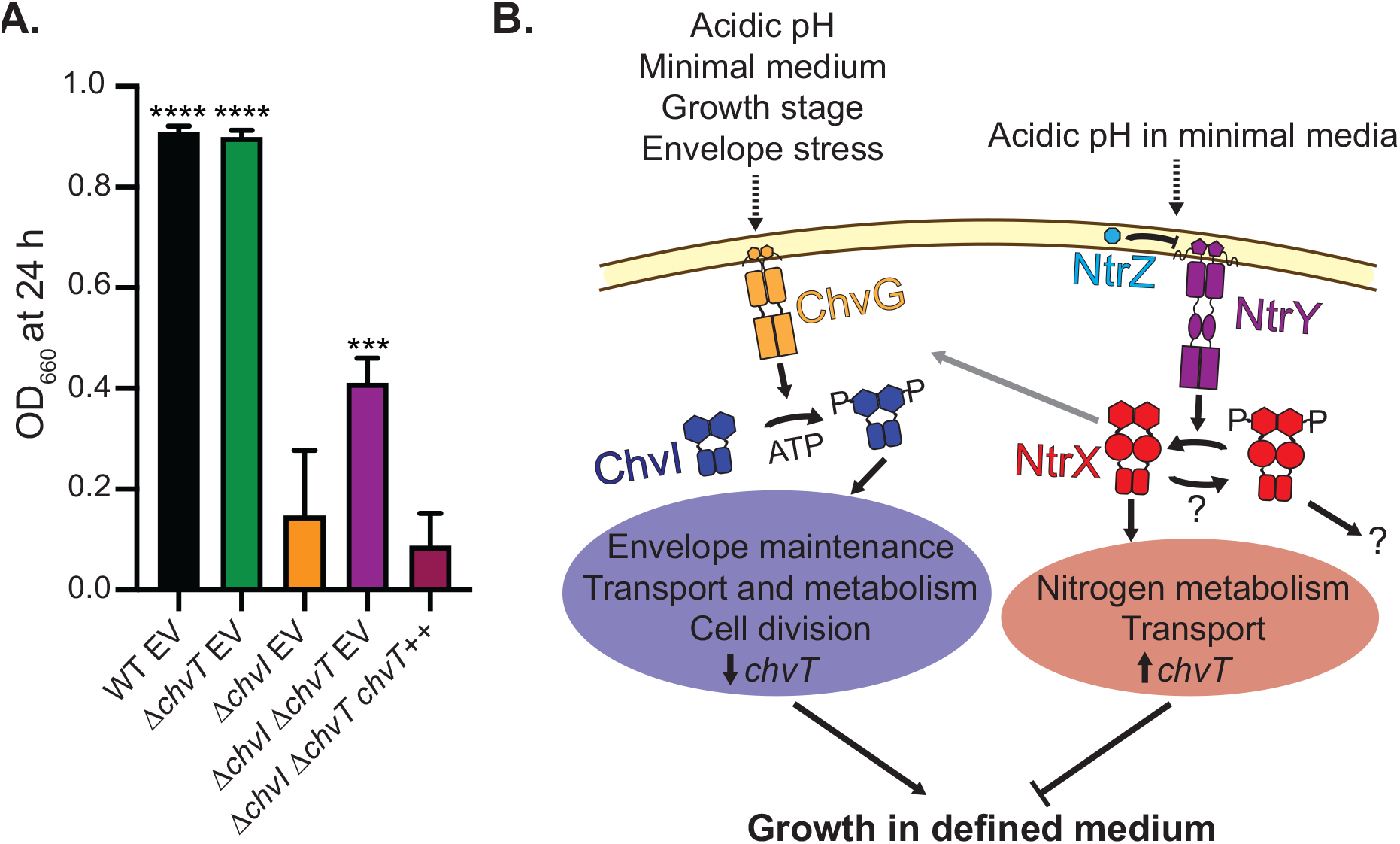
Deletion of *chvT* partially restores growth of *ΔchvI* cells in M2X medium. (A) Optical density (OD_660_) of WT, ΔchvT, and *ΔchvI ΔchvT* strains, with empty vector (EV) or *chvT* overexpression vector (++), 24 h after back-dilution to OD_660_ = 0.025 in M2X medium (average ± SD, *N* = 4). ^***^ = *p* < 0.0005, ^****^ = *p* < 0.0001, one-way ANOVA followed by Dunnett’s post-test comparison to *ΔchvI* EV. (B) Proposed model for the regulatory interactions between ChvGI and NtrYXZ. ChvG (orange) is activated by a variety of cellular conditions (dashed arrow) and phosphorylates ChvI (blue). Phosphorylated ChvI regulates genes involved in envelope maintenance, transport, metabolism, and cell division (blue oval). In addition, ChvI represses expression of *chvT*. NtrY (purple) is repressed by NtrZ under acidic conditions in defined medium (dashed arrow), reducing dephosphorylation of NtrX (red). Unphosphorylated NtrX regulates much of the ChvI regulon, likely via upregulation of the *chvIG-hprK* operon (grey arrow). In addition, unphosphorylated NtrX upregulates *chvT* and regulates expression of genes involved in nitrogen metabolism and transport (red oval). Phosphorylated ChvI and unphosphorylated NtrX oppose each other in regulating growth in defined medium, partially via differential regulation of *chvT*. The transcriptional role of phosphorylated NtrX and source of phosphorylation (question marks) are not yet known.

## Discussion

### The importance of the *C. crescentus* ChvGI system for growth in defined medium

To our knowledge, the only reported physiological phenotype for *chvGI* mutants in *C. crescentus* is sensitivity to the antibiotic vancomycin (47). However, previous work indicated that ChvGI might be important for growth in defined medium (6). In fact, *chvGI* mutants in other α-proteobacteria are sensitive to nutritional conditions, although most, with the exception of *Brucella abortus*, exhibit particularly poor growth in complex media (4, 8, 27, 59). Deletion of *chvG* or *chvI* in *C. crescentus* caused a distinctive growth defect in M2X medium. *ΔchvI* cultures do grow in M2X upon inoculation from PYE plates, suggesting that the physiological state of the cell in PYE agar is initially amenable to growth in defined medium (Fig. 1). However, this tolerance is limited by time, with only higher inocula able to reach high cell density (Fig. 1 and Fig. S2). Why then, can primary overnight cultures persist at high density until back-dilution in fresh M2X? One possibility is that the low pH of M2X at stationary phase preserves *ΔchvI* and *ΔchvG* cells, potentially by triggering phosphorylation of NtrX (Fig. 7B) (48). This model is supported by the observation that *ΔchvI* cells reach higher CFUs when back-diluted in M2X at pH 5.5 vs. pH 6.0 (Fig. S4C). As ∼20% of the NtrX pool is phosphorylated in M2G medium at pH 6.0 vs. ∼40% at pH 5.5 (by Phos-tag analysis), the pH 6–5.5 transition would significantly change the level of unphosphorylated NtrX (48). However, as *ΔchvI* and *ΔchvG* cells also appear to be sensitive to acidic pH, additional factors are likely at play (Fig. S4C). For example, several extracytoplasmic function (ECF) sigma factors are involved in resistance to stationary-phase stress and might play a protective role in *ΔchvI* and *ΔchvG* cells (60-62).

A recent study failed to identify *chvG* or *chvI* as being important for fitness in M2X medium (63). However, in this work M2X cultures were inoculated from PYE starter cultures at high enough density to reach saturation in 5 doublings. Thus, these experiments likely mimicked our primary overnight cultures, obscuring detection of fitness defects for *chvGI* mutants.

### The ChvI transcriptional regulon in *C. crescentus*

Although ExoR or ChvI transcriptional regulons have been defined in several α-proteobacteria (4, 5, 26, 64), the *C. crescentus* ChvI regulon was unknown prior to this work. We employed RNA-seq to detect direct and indirect transcriptional targets of ChvI in *C. crescentus* (Fig. 6, Table S1). Our ChvI regulon contained several classes of genes noted in other α-proteobacteria, including those encoding outer membrane proteins and transporters, metabolic enzymes, lipopolysaccharide biosynthesis enzymes, and stress response proteins (4, 5, 26, 64). We note, in particular, that the *C. crescentus* ChvI regulon contained a large number of genes encoding proteins involved in envelope maintenance, including nearly the entire BAM complex, the envelope integrity protein EipA, members of the Tol-Pal complex, and a variety of chaperones and proteases (Fig. 6 and Table S1) (65-68). The idea that ChvGI is involved in envelope integrity is consistent with previous observations that ChvGI is activated by envelope stress and confers resistance to antibiotics targeting the cell wall (47). However, ∼40% of genes regulated by ChvI are annotated generically or as hypotheticals, and thus, more work will be required to characterize the pathways downstream of the ChvGI system. In *A. tumefaciens, S. meliloti*, and *C. crescentus*, ChvGI appears to suppress cell motility and/or chemotaxis (5, 15, 20, 26, 69, 70). However, unlike in *A. tumefaciens* and *S. meliloti, C. crescentus* ChvI does not regulate any obvious flagellar or chemotaxis genes, suggesting that effects on motility may be due to post-transcriptional regulation and/or alterations in cell-cycle progression (5, 26, 71-73).

Overexpression of the phosphomimetic *chvI(D52E)* allele induced cell filamentation in M2X, implicating ChvI in regulating cell division, and cytokinesis in particular. ChvI upregulates several genes related to cell division, including *zauP*, members of the Tol-Pal complex, *smc*, and *nstA* (68, 74-77). We note that overexpression of a proteolytically stable mutant form of *nstA* induces cell filamentation and both *zauP* and the Tol-Pal complex are involved in regulating cytokinesis (74, 77). Perhaps over-induction of these regulon genes, in combination with the cellular state in M2X medium, interferes with proper cell division.

### Genetic interactions between ChvGI and NtrYX

The severe growth defect of *ΔchvG* and *ΔchvI* cells in M2X medium allowed us to uncover the first known genetic interaction between *chvGI* and *ntrYX* (Fig. 3). Importantly, deletion of *ntrX* suppressed the growth defect of *ΔchvI* cells, whereas overexpression of non-phosphorylatable *ntrX(D53A)* was deleterious for growth in both *ΔchvI* and WT cells (Fig. 4). These results suggest that the activity of unphosphorylated NtrX is particularly detrimental in cells lacking *chvI*. Although unphosphorylated response regulators are often assumed to be inactive, multiple RRs are known to affect transcription in their unphosphorylated states (78-82). Thus, we propose that phosphorylated ChvI and unphosphorylated NtrX oppose each other to regulate growth in defined medium (Fig. 7B). We note that a past study examining connections between ChvI and NtrX in *S. meliloti* did not test whether perturbations in NtrYX signaling might affect *ΔchvI* phenotypes (25). We predict that a similar ChvI–NtrX relationship may be conserved in other α-proteobacteria.

As gain-of-function mutations in *ntrY* and *ntrZ* suppress the growth defect of *ΔchvI* cells, we also propose that phosphorylation of NtrX relieves its detrimental activity, possibly by changing its transcriptional regulon. Although the global transcriptional effects of NtrX phosphorylation have yet to be characterized, *in vitro* phosphorylation of *B. abortus* NtrX does induce conformational changes and alters, but does not weaken, binding to the *ntrYX* promoter (83). Interestingly, we were unable to construct *ΔchvI ΔntrY* and *ΔchvI ΔntrZ* strains, suggesting that these gene deletion combinations are synthetically lethal in PYE. As *ntrX* is also essential in PYE (20), the balance between unphosphorylated and phosphorylated NtrX may be important for growth in both complex and defined media.

To identify downstream genes that mediate the oppositional relationship between ChvI and NtrX, we compared their transcriptional regulons (Fig. 6 and Table S1). The ChvI regulon strongly overlapped with that of NtrX, as 80% of the top 30 ChvI-activated genes are also activated by NtrX (per our >1.5-fold cutoff). By contrast, only 20% of the top 30 NtrX-activated genes exhibit >1.5-fold ChvI-dependence (49). Given that *chvI, chvG*, and *hprK* are upregulated by NtrX, it may be the case that NtrX simply activates expression of the *chvIG-hprK* operon, thereby altering transcription of genes in the ChvI regulon (Fig. 7B, grey arrow). However, further work is required to define the mechanism by which NtrX affects the ChvI regulon, as it may also indirectly affect ChvGI signaling or directly regulate expression of regulon genes. The overlap we observed between the ChvI and NtrX regulons may be restricted to *C. crescentus* and close relatives, as *chvG* and *chvI* are not transcriptionally regulated by NtrYX in *Rhodobacter sphaeroides* (40). However, ChvGI and NtrYX may still be transcriptionally linked in more distantly-related α-proteobacterial species, as *ntrX* is part of the *A. tumefaciens* ExoR regulon (26).

The majority of overlapping genes in the ChvI and NtrX regulons cannot account for the detrimental effect of unphosphorylated NtrX on *ΔchvI* cells. Therefore, we focused on eight genes that exhibited opposing transcriptional regulation by ChvI and NtrX (Fig. S6). Given that NtrX largely reinforces the ChvGI TCS, these oppositional effects may reflect direct transcriptional regulation by NtrX or the effects of unique NtrX regulon genes. Only deletion of *chvT* improved the growth of *ΔchvI* cells, indicating that suppression of *chvT* RNA levels by ChvI may be important for growth in M2X medium (Fig. 7). *chvT* is linked to diverse phenotypes in *C. crescentus*, including survival in stationary phase, sensitivity to cell wall-targeting antibiotics, and sensitivity to bacteriocins (47, 84, 85). However, the molecule(s) transported by ChvT remain undefined. High ChvT levels may contribute to defects in transport in defined medium and/or alter membrane integrity, impacting the viability of cells lacking ChvGI (47). It is clear, though, that altered *chvT* expression cannot fully explain the growth deficiency of *ΔchvI* cells in M2X medium. This growth defect may result from the collective action of several genes, and therefore, manipulation of individual candidates may not rescue growth in M2X. Moreover, 15 genes in the ChvI regulon are absent from the NtrX microarray dataset, raising the possibility that one or more may be differentially regulated by ChvI and NtrX (Fig. 6, Table S1). ChvI and NtrX might also interact more indirectly, as each regulates unique subsets of genes that may affect growth in M2X medium.

### On the role of NtrZ

NtrYX is associated with a wide range of physiological responses, from nitrogen metabolism to redox sensing and cell envelope maintenance (25, 35, 39, 40). Despite these phenotypic observations, little is known about NtrY activity *in vivo* and the regulation of NtrY via its periplasmic domain. Our work reveals a surprising phosphatase pathway involving the previously uncharacterized protein NtrZ. Phos-tag analysis of NtrX phosphorylation *in vivo* clearly demonstrates that NtrY is dispensable for NtrX phosphorylation, and suggests that NtrY primarily acts as an NtrX phosphatase (Fig. 5). In addition, NtrZ appears to inhibit NtrY phosphatase activity, as deletion of *ntrZ* abolishes NtrX phosphorylation only when *ntrY* is present (Fig. 5). Thus, we propose that NtrZ inhibits NtrY phosphatase activity, stabilizing the pool of phosphorylated NtrX (Fig. 7). In the future, we are interested in determining whether NtrZ physically interacts with the NtrY periplasmic domain or affects NtrY activity indirectly. Several known periplasmic or membrane-bound TCS regulators directly interact with HK periplasmic domains (33, 86-88). Although our model does not exclude the possibility that NtrY phosphorylates NtrX under certain conditions, there is clearly another source of NtrX phosphorylation in *C. crescentus*. An additional HK may phosphorylate NtrX, although the most likely candidate, NtrB, does not phosphorylate NtrX *in vitro* (49). Alternatively, some metabolic intermediates, such as aspartyl-phosphate or carbamoyl-phosphate, can serve as phosphodonors for RRs *in vivo* and *in vitro* (1, 89-92).

Interestingly, we initially placed *ntrZ* upstream of *ntrY* by examining the similar stationary-phase survival phenotypes of *ΔntrY* and *ΔntrZ* cells (Fig. 5). Given our Phos-tag results, these phenotypes suggest that both phosphorylated and unphosphorylated NtrX play important roles in stationary-phase survival. This idea is consistent with the fact that NtrX phosphorylation increases upon entry into stationary-phase, but eventually decreases (48).

Fernández et al. reported that NtrX phosphorylation is triggered by acidic pH in defined medium (48). Therefore, we hypothesize that inhibition of NtrY by NtrZ may be enhanced by acidic pH. Thus, NtrZ and ExoR potentially share a regulatory theme, in which their activities towards their respective HKs are affected by low pH (16, 33, 34). However, it remains to be seen if, as with ExoR, proteolysis plays a role in controlling NtrZ-dependent regulation of NtrY (16). Given the results of our suppressor selection, it would appear that mutations in NtrZ or in the transmembrane helices of NtrY may bypass regulation by acidic pH. In an alignment of 100 NtrZ homologs, ranging from 41% to 100% sequence identity, the Y92 position is conserved as an aromatic residue (95% Y) while I99 is conserved as an aliphatic hydrophobic residue (51% I, common L and V substitutions), suggesting that these residues are important for NtrZ function. Further work is required to determine how mutation of Y92 or I99 may alter interaction with the NtrY periplasmic domain or affect a different aspect of NtrZ function. An alignment of the non-cytoplasmic portion of 250 NtrY sequences, ranging from 37% to 100% identity, reveals that the L70 position is largely conserved as an aliphatic hydrophobic residue (94% L) whereas A123 is conserved as a hydrophobic residue (93% A), albeit with occasional bulky-aromatic substitutions. It is unsurprising, then, that the L70H and A123V substitutions appear to affect NtrY function. In fact, several studies have identified transmembrane mutations that alter HK kinase and phosphatase activities (93-95).

Unlike the periplasmic kinase regulator ExoR, which contains Sel1-like repeats typically involved in protein-protein interactions, NtrZ does not contain any conserved domains or motifs (95). Moreover, NtrZ appears restricted to the order Caulobacterales, albeit with a few distant homologs in other α-proteobacteria. Notably, neither *B. abortus, S. meliloti*, nor *A. tumefaciens* contain NtrZ homologs, and conservation of the non-cytoplasmic region of NtrY is quite low between *C. crescentus* and these organisms (24–27% identity). However, L70 is largely conserved (L or V) and A123 is entirely conserved between these NtrY homologs. Thus, the L70H and A123V substitutions may have similar effects on NtrY activity in these organisms, and further analyses of these mutants may shed light on a conserved NtrY activity switch. Moreover, it will be interesting to see if NtrY also primarily acts as a phosphatase and/or is regulated by periplasmic effectors in *S. meliloti, A. tumefaciens*, and *B. abortus*.

## Materials and Methods

### Strains and Plasmids

All plasmids were constructed using standard molecular biology techniques. See Table S2 for strain, plasmid, and primer information. Plasmids for generating in-frame deletions and allele-replacements were generated by cloning homologous upstream and downstream regions into pNPTS138. Transcriptional reporter plasmids were generated by cloning 400–500 bp upstream of the open reading frame into pRKlac290. For overexpression strains, ORFs were inserted into pMT585, a plasmid for xylose-inducible expression that integrates at the *xylX* locus. Plasmids were transformed into *C. crescentus* CB15 strain by electroporation or tri-parental mating. In-frame deletion and allele-replacement strains were generated by a double recombination strategy involving *sacB* counterselection on PYE plates supplemented with 3% sucrose (97). For *ΔntrX* knockout strains, counterselection was carried out on M2X + 0.5% sucrose plates due to their growth defect on PYE. *ΔntrX* and *ΔchvI ΔntrX* strains were grown only on M2 medium. Construction of *ΔchvI ΔntrY* and *ΔchvI ΔntrZ* mutants was attempted using both PYE and M2X counterselection methods. All *C. crescentus* strains were grown at 30°C. For strains carrying pRKlac290 plasmids, oxytetracycline was added to 1 µg/mL in liquid and 2 µg/mL in PYE agar or 1 µg/mL in M2X agar.

M2X medium contained 6.1 mM Na_2_HPO_4_, 3.9 mM KH_2_PO_4_, 9.3 mM NH_4_Cl, 0.25 mM CaCl_2_, 0.5 mM MgSO_4_, 1:1000 100X Ferrous Sulphate/Chelate Solution (Sigma), and 0.15% xylose. PYE medium contained 0.2% peptone, 0.1% yeast extract, 0.5 mM CaCl_2_, and 1 mM MgSO_4._

### Measurement of Growth in M2X Medium

Primary M2X cultures were inoculated from plates to an approximate density of OD_660_ 0.02-0.10 and grown overnight. Overnight cultures were back-diluted to OD_660_ = 0.025 and OD_660_ was recorded at the indicated times. To enumerate colony forming units, samples were taken at the indicated time points and 10-fold serial dilutions were plated on PYE agar. For pH experiments, M2X medium was adjusted to the indicated pH using HCl.

For experiments with controlled starting densities, cells were resuspended in M2X medium directly from PYE plates. These resuspensions were then diluted to the indicated OD_660_ and the cultures were titered for CFUs. In washing experiments, 1 mL resuspended cells were spun down at 8k × *g* for 3 min and resuspended in 1 mL fresh M2X medium once (1X) or twice (2X) before dilution. Plotting and statistical analyses were carried out using Prism (Graphpad).

### Microscopy

Samples of *ΔchvG* and *ΔchvI* cells were taken from overnight M2X or PYE + 0.15% xylose cultures and imaged with a DMI6000B (Leica) microscope in phase contrast using a HC PL APO 63x/1.4na oil Ph3 CS2 objective. Images were captured using an Orca-R^2^ C10600 digital camera (Hamamatsu) controlled by Leica Application Suite X (Leica). Images were processed using Fiji (98, 99).

### Suppressor Screen

Overnight M2X cultures of *ΔchvG* and *ΔchvI* cells were back-diluted to OD_660_ = 0.025 in M2X medium. Cultures initially saturated at OD_660_ ∼0.1 before growing to higher density (OD_660_ = 0.5– 0.8) after 2-3 days growth. From each culture, single colonies were isolated on PYE plates. Isolated strains were then tested for suppression by M2X growth curves. The origin of each suppressor strain is as follows: culture 1 (*ΔIS1*), culture 2 (*ΔIS2, ΔIS3*), culture 3 (*ΔGS1*), culture 4 (*ΔGS2, ΔGS3*). Genomic DNA was isolated from 1 mL of overnight culture grown in PYE medium, using guanidinium thiocyanate (100). Sequencing was performed by the Microbial Genome Sequencing Center (Pittsburgh, PA) using a single library preparation method based upon the Illumina Nextera Kit. Sequences were aligned to the *C. crescentus* NA1000 reference genome (GenBank accession number CP001340) using breseq (101).

### Phos-tag gel electrophoresis and western blotting

2 mL M2X overnights of strains containing *ntrX-HA* at the native *ntrX* locus were grown and back-diluted to OD_660_= 0.01. Samples were collected after 22 h (0.25 mL•OD_660_; i.e. volume [mL] = 0.25/OD_660_ of culture) and frozen at -80°C. Samples were thawed, resuspended in 2.5X SDS loading buffer (125 mM Tris [pH 6.8], 25% glycerol, 5% SDS, 5 mM DTT, 0.01% bromophenol blue) containing 1:50 benzonase (Sigma), and immediately loaded onto Phos-tag gels.

Phos-tag electrophoresis was performed as described (48) using 8% acrylamide gels copolymerized with 35 µM Phos-tag acrylamide (NARD) and 150 µM ZnCl_2_. Proteins were transferred to polyvinylidene difluoride (PVDF; Bio-Rad) in a wet-transfer apparatus. Membranes were probed with monoclonal HA Tag antibody (1:2000 dilution, Invitrogen, 2-2.2.14,), incubated with anti-mouse IgG-horseradish peroxidase (IgG-HRP) conjugate (1:5000 dilution, Invitrogen), and developed with ProSignal™ Pico Spray (Prometheus). Blots were imaged using a Bio-Rad ChemiDoc MP Imager. Bands were quantified using Fiji (98, 99).

### Transcriptome Deep Sequencing (RNA-seq)

2 mL of PYE medium was inoculated with *ΔchvI* EV (*ΔchvI xylX*::pMT585) and *ΔchvI chvI(D52E)++* (*ΔchvI xylX*::pMT585-*chvI(D52E)*) cells and grown overnight. Cultures were diluted to OD_660_ = 0.001 in 2 mL fresh PYE and grown for 22.5 h. Cultures were diluted to OD_660_ = 0.075 in 5 mL PYE + 0.15% xylose and grown for 3.5 h before TRIzol extraction and RNA isolation, as described previously (102). RNA-seq libraries were prepared using an Illumina TruSeq stranded RNA kit and sequenced on an Illumina NextSEQ500 instrument at the University of Chicago Functional Genomics Facility. Sequencing data were analyzed using CLC Genomics Workbench 20 (Qiagen) by mapping reads to the *C. crescentus* NA1000 genome (84). Motif searching was carried out using MEME (103). Heatmaps were generated using Java Treeview3 and hypergeometric analysis was performed using phyper in R (104, 105).

### β-galactosidase Assays

For assays of WT vs. *ΔchvI* strains, cells were grown overnight in 2 mL PYE medium. Then, 500 µL of each culture was centrifuged at 8k × *g* for 3 min and resuspended in 500 µL M2X medium. Resuspended cultures were used to inoculate 2 mL M2X medium at a starting OD_660_ of 0.075. Cultures were grown for 4 h before assaying β-galactosidase activity. For assays of WT vs. *ΔntrX* strains, cells were grown overnight in 2 mL M2X medium. Overnight cultures were diluted in M2X medium such that they would reach OD_660_= 0.1–0.2 after 23.5 h of growth. β-galactosidase assays were performed as previously described using 200 µL of M2X culture + 100 µL sterile PYE medium as an emulsifier (106). Plotting and statistical analyses were carried out using Prism (Graphpad).

## Data Availability

RNA-seq data are available in the NCBI’s Gene Expression Omnibus (GEO) database (107) under the GEP Series accession number GSE168965 (https://www.ncbi.nlm.nih.gov/geo/query/acc.cgi?acc=GSE168965).

## Acknowledgements

We thank Clare Kirkpatrick, Régis Hallez, Alex Quintero, and members of the Crosson laboratory for helpful discussion, and Jen Mach and Plant Editors for constructive feedback on the manuscript. Research in this publication was supported by the National Institute of General Medical Sciences of the National Institutes of Health (NIH) under Award Numbers F32 GM128283 (B.J.S.) and R35 GM131762 (S.C.). The content is solely the responsibility of the authors and does not necessarily represent the official views of the National Institutes of Health.

