## Supplemental Materials for "The ChvG–ChvI and NtrY–NtrX two-component systems coordinately regulate growth of *Caulobacter crescentus*"

##### Supplemental Figures

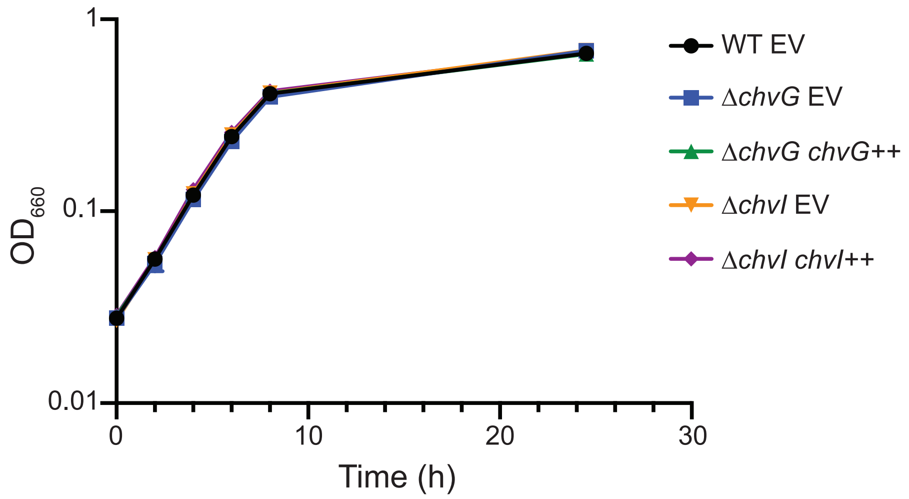


**Figure S1: Deletion of *chvI* and *chvG* does not affect growth in PYE medium**
Growth curves, measured by optical density ($OD_{660}$), of WT and knockout strains upon back-dilution in PYE medium. Strains bear empty vector (EV) or genetic rescue plasmids (++).

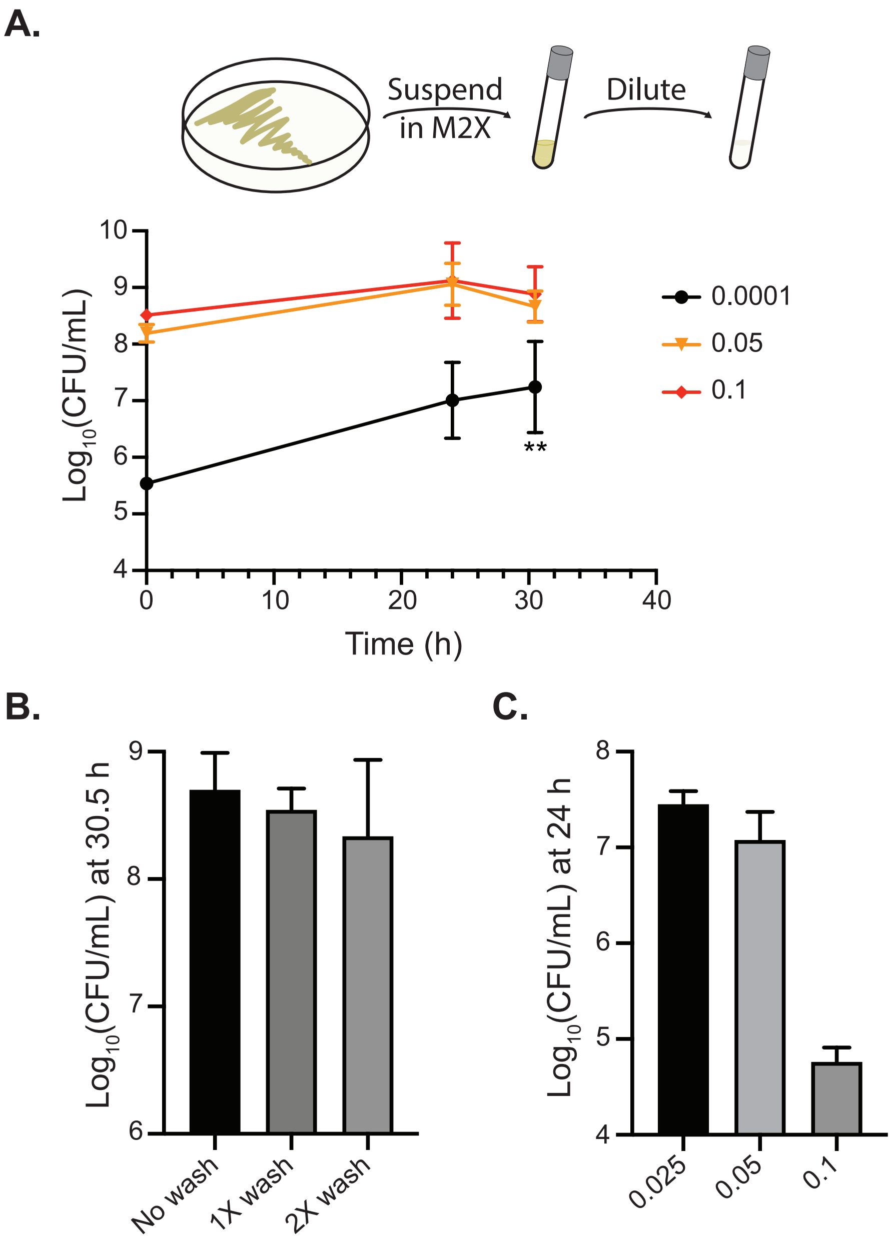


**Figure S2: Starting culture density, but not PYE contamination, determines whether Δ*chvI* overnight cultures reach high density in M2X medium.**

(A) Growth measured by CFU for Δ*chvI* cultures diluted to the indicated starting $OD_{660}$ in M2X medium. Points represent averages of four biological replicates ± SD. ** = $p < 0.01$, one-way ANOVA followed by Dunnett's post-test comparison to the culture starting at $OD_{660} = 0.1$. (B) $Log_{10}$(CFU/mL) values for Δ*chvI* cultures started at a density of $OD_{660} = 0.05$ and grown for 30.5 hours in M2X medium (average ± SD, $N = 3$). Before dilution to the starting density, cells were resuspended in 1 mL M2X medium from a PYE plate (No wash) and washed once (1X wash) or twice (2X wash) with 1 mL M2X medium. (C) $Log_{10}$(CFU/mL) values for Δ*chvI* cultures back-diluted from overnight cultures in M2X medium to the indicated starting $OD_{660}$ and grown for 24 h (average ± SD, $N = 3$).

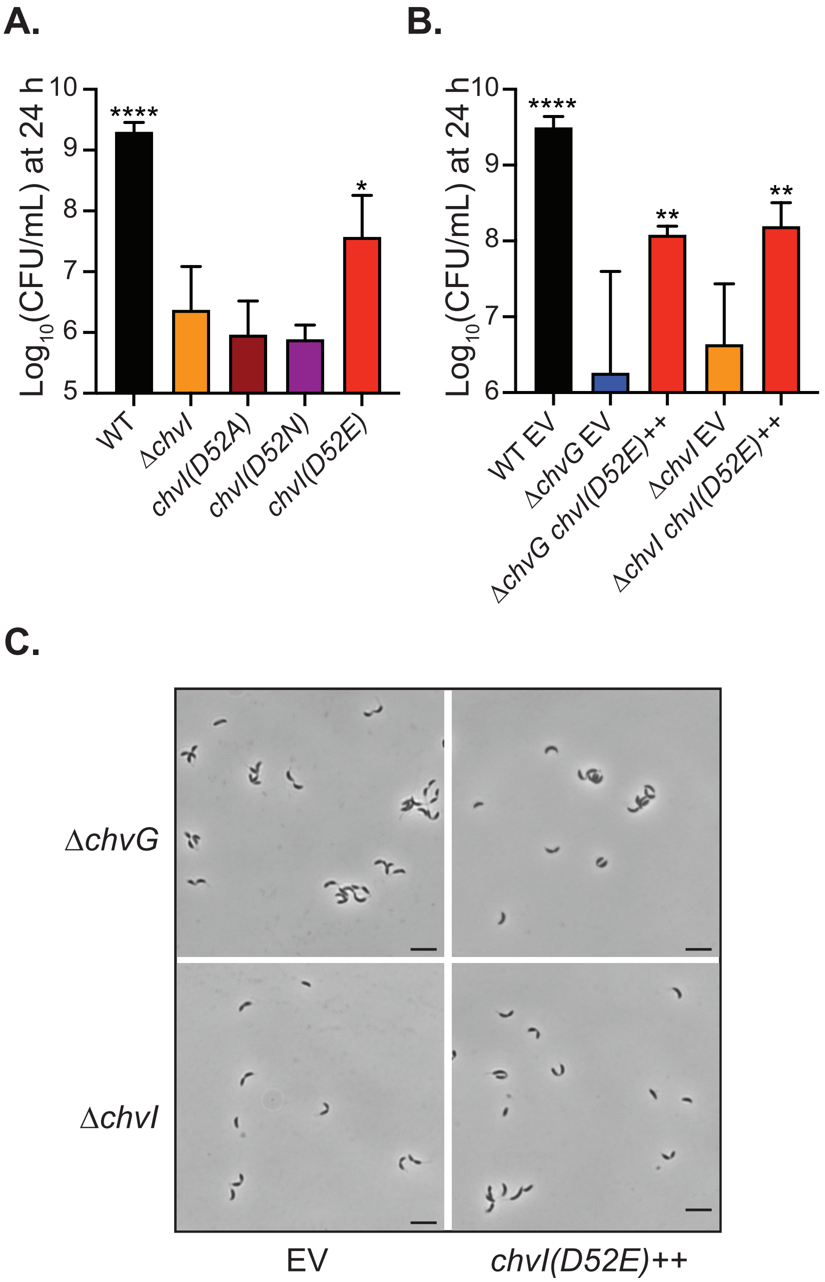


**Figure S3: Viable cell counts are consistent with $OD_{660}$ measurements for *chvI* allele replacement and overexpression strains.**

(A) $Log_{10}$(CFU/mL) values for WT and *chvI* mutant strains 24 h after back-dilution in M2X medium (average ± SD, $N \geq 4$). * = $p < 0.05$, **** = $p < 0.0001$, one-way ANOVA followed by Dunnett's post-test comparison to Δ*chvI*. **(**B) $Log_{10}$(CFU/mL) values for WT and mutant strains, bearing empty vector (EV) or overexpression vectors (++), 24 hours after back-dilution in M2X medium (average ± SD, $N = 3$). * = $p < 0.05$, ** = $p < 0.01$, **** = $p < 0.0001$, one-way ANOVA followed by Dunnett's post-test comparison to Δ*chvG*. WT, Δ*chvG* EV, and Δ*chvI* EV data are from Fig. 1B. (C) Phase contrast micrographs of primary overnight cultures grown in PYE medium + 0.15% xylose. Strains carry empty vector (EV) or a *chvI(D52E)* overexpression vector. Scale bars, 5 µm.

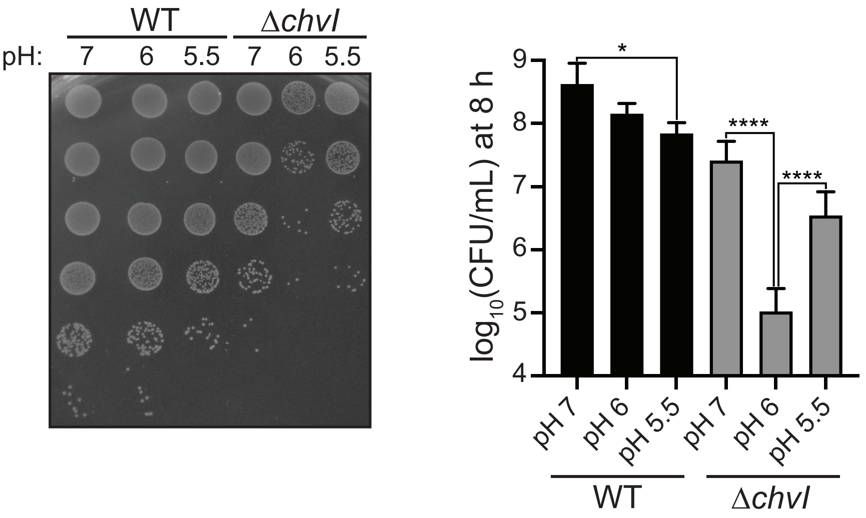


**Figure S4: pH has a bimodal effect on growth of Δ*chvI* cells in M2X medium.**
$Log_{10}$(CFU/mL) values for WT and Δ*chvI* strains 8 h after back-dilution in M2X medium at the indicated pH. Representative serial dilutions (left) and quantification (right, average ± SD, $N = 4$). * = $p < 0.05$, **** = $p < 0.0001$, one-way ANOVA followed by Tukey's post-test comparison.

**A.**

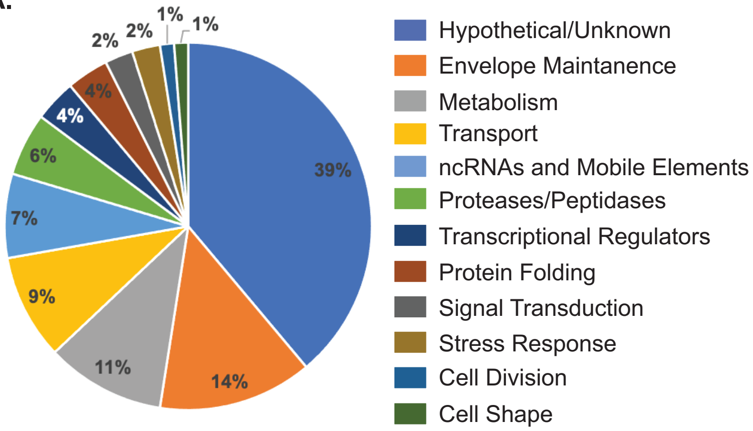


**B.**

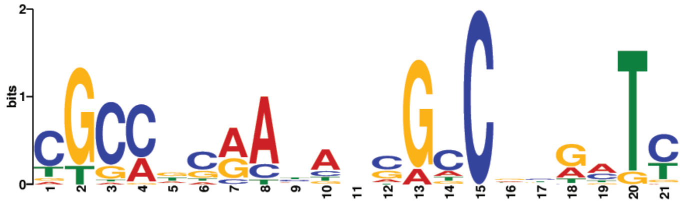


**Figure S5**: **Characteristics of the ChvI transcriptional regulon**
(A) Pie chart of ChvI regulon genes organized into functional categories. Genes were manually classified based upon annotations and published data. (B) Sequence logo of the putative ChvI binding site in *C. crescentus*. Results of MEME (1) search using 100 bp upstream of the experimentally-determined transcription start site (TSS) (2) or 200 bp upstream of the start codon for the top 35 upregulated operons. This motif was identified in 30/35 sequences with an overall E-value of $2.8\times10^{-9}$. The 3' end of motif was located ≤ 30 bp upstream of the TSS in 10/11 operons with known TSSs.

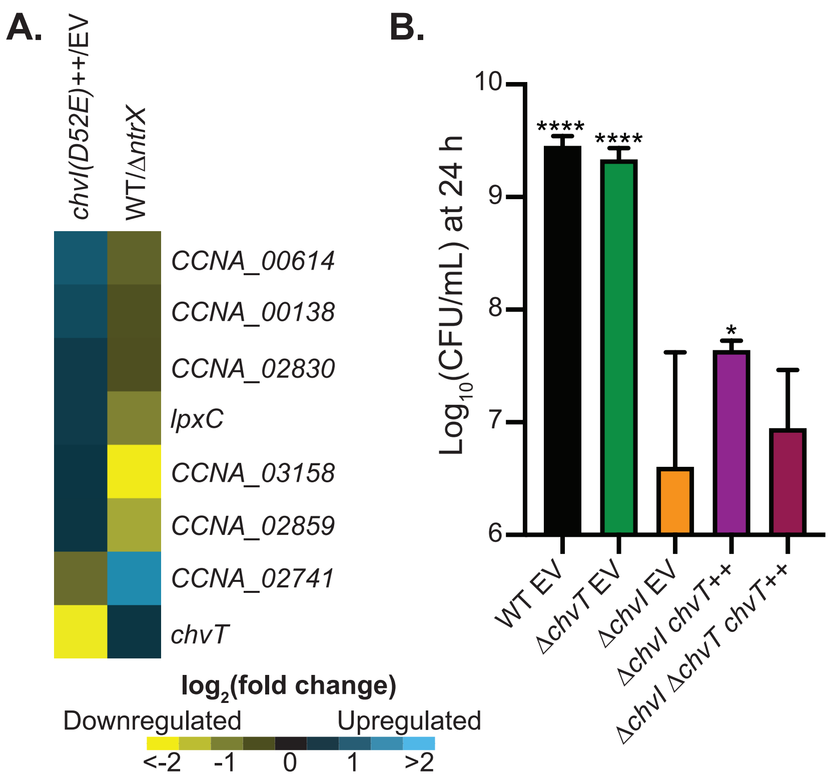


**Figure S6: Differential regulation of *chvT* by ChvI and NtrX may affect growth in M2X medium.**
(A) Subset of the heatmap in Fig. 6A showing genes that are differentially regulated by ChvI and NtrX. (B) $Log_{10}$(CFU/mL) values for WT, Δ*chvT*, and Δ*chvI* Δ*chvT* strains, with either empty vector (EV) or the *chvT* overexpression vector (++), 24 h after back-dilution in M2X medium (average ± SD, $N = 4$). * = $p < 0.05$, **** = $p < 0.0001$, one-way ANOVA followed by Dunnett's post-test comparison to Δ*chvI* EV.

#### Table S1: The ChvI regulon

| Locus | Annotation | Max group mean (RPKM) | Fold Change (Δ*chvI chvI(D52E)*++/Δ*chvI* EV) | $Log_2$ (Fold Change) | *p*-value | FDR *p*-value | Fold Change (WT/Δ*ntrX*)[3] |
|---|---|---|---|---|---|---|---|
| ***CCNA_R0092*** | minimal medium expressed sRNA | 4.56 | 182.15 | 7.51 | 7.99E-06 | 1.12E-04 | |
| ***CCNA_03987*** | hypothetical protein | 377.22 | 128.67 | 7.01 | 0.00E+00 | 0.00E+00 | |
| ***chvR*** | small non-coding RNA | 9.22 | 70.97 | 6.15 | 5.57E-12 | 1.43E-10 | |
| ***CCNA_02817*** | conserved hypothetical protein | 10.59 | 56.41 | 5.82 | 0.00E+00 | 0.00E+00 | 1.37 |
| ***CCNA_R0088*** | minimal medium expressed sRNA | 6.60 | 48.18 | 5.59 | 3.22E-15 | 9.98E-14 | |
| ***CCNA_00733*** | GumN superfamily protein | 9.42 | 47.20 | 5.56 | 0.00E+00 | 0.00E+00 | 5.48 |
| ***CCNA_00889*** | NTF2-family protein | 157.29 | 45.18 | 5.50 | 0.00E+00 | 0.00E+00 | 4.39 |
| ***CCNA_03997*** | amelogenin/CpxP-related protein | 27.74 | 33.24 | 5.05 | 0.00E+00 | 0.00E+00 | |
| ***CCNA_01238*** | EF-hand domain protein | 24.22 | 26.67 | 4.74 | 0.00E+00 | 0.00E+00 | 4.04 |
| ***CCNA_02009*** | hypothetical protein | 12.24 | 26.60 | 4.73 | 0.00E+00 | 0.00E+00 | 2.94 |
| ***chvG*** | two-component sensor histidine kinase chvG | 43.72 | 16.00 | 4.00 | 0.00E+00 | 0.00E+00 | 5.03 |
| ***CCNA_02531*** | proline-rich hypothetical protein | 1.51 | 15.94 | 3.99 | 0.00E+00 | 0.00E+00 | 2.91 |
| ***CCNA_00546*** | conserved hypothetical protein | 13.08 | 15.04 | 3.91 | 0.00E+00 | 0.00E+00 | 4.18 |
| ***CCNA_03308*** | hypothetical protein | 19.10 | 14.56 | 3.86 | 0.00E+00 | 0.00E+00 | 3.27 |
| ***CCNA_01090*** | conserved hypothetical protein | 93.68 | 14.34 | 3.84 | 0.00E+00 | 0.00E+00 | 2.80 |
| ***CCNA_01660*** | conserved hypothetical protein | 46.55 | 13.39 | 3.74 | 0.00E+00 | 0.00E+00 | 3.25 |
| ***CCNA_01443*** | YiijF-family protein | 67.30 | 13.32 | 3.74 | 0.00E+00 | 0.00E+00 | 11.66 |
| ***CCNA_01341*** | DegP/HtrA-family serine protease | 38.48 | 11.53 | 3.53 | 0.00E+00 | 0.00E+00 | 2.34 |
| ***CCNA_02309*** | EF hand domain protein | 37.08 | 11.39 | 3.51 | 0.00E+00 | 0.00E+00 | 6.38 |
| ***hprK*** | serine kinase/phosphatase HprK | 3.80 | 9.38 | 3.23 | 0.00E+00 | 0.00E+00 | 3.90 |
| ***CCNA_01089*** | conserved hypothetical protein | 19.62 | 9.35 | 3.23 | 0.00E+00 | 0.00E+00 | 4.64 |
| ***CCNA_02202*** | YkwD-family protein | 5.09 | 9.20 | 3.20 | 0.00E+00 | 0.00E+00 | 2.10 |
| ***CCNA_R0095*** | small non-coding RNA | 1.62 | 7.80 | 2.96 | 0.00E+00 | 0.00E+00 | |
| ***CCNA_02339*** | class I SAM-dependent methyltransferase | 9.51 | 6.30 | 2.66 | 0.00E+00 | 0.00E+00 | 1.31 |
| ***CCNA_01505*** | MreB-interacting protein MbiA | 30.66 | 5.63 | 2.49 | 0.00E+00 | 0.00E+00 | 2.61 |
| ***CCNA_02001*** | HAD/PGPPase-family phosphohydrolase | 5.96 | 5.44 | 2.44 | 0.00E+00 | 0.00E+00 | 1.58 |

| | | | | | | | |
|---|---|---|---|---|---|---|---|
| ***CCNA_02338*** | hypothetical protein | 20.71 | 5.02 | 2.33 | 0.00E+00 | 0.00E+00 | 1.39 |
| ***CCNA_R0106*** | small non-coding RNA | 89.65 | 4.96 | 2.31 | 0.00E+00 | 0.00E+00 | |
| ***CCNA_00038*** | hypothetical protein | 21.53 | 4.86 | 2.28 | 0.00E+00 | 0.00E+00 | 2.05 |
| ***CCNA_03820*** | LolA-family outer membrane lipoprotein carrier protein | 127.08 | 4.81 | 2.27 | 0.00E+00 | 0.00E+00 | 1.42 |
| ***CCNA_00039*** | hypothetical protein | 44.46 | 4.66 | 2.22 | 0.00E+00 | 0.00E+00 | 1.99 |
| ***eipA*** | cell envelope integrity protein EipA | 424.34 | 4.48 | 2.16 | 0.00E+00 | 0.00E+00 | |
| ***CCNA_01080*** | hypothetical protein | 33.43 | 4.32 | 2.11 | 0.00E+00 | 0.00E+00 | 1.61 |
| ***CCNA_03439*** | hypothetical protein | 11.47 | 4.13 | 2.04 | 0.00E+00 | 0.00E+00 | 1.02 |
| ***CCNA_02165*** | lysophospholipase | 3.96 | 3.93 | 1.97 | 0.00E+00 | 0.00E+00 | 2.00 |
| ***CCNA_00484*** | hypothetical protein | 20.27 | 3.90 | 1.96 | 0.00E+00 | 0.00E+00 | 3.34 |
| ***CCNA_02815*** | ice nucleation protein | 15.30 | 3.73 | 1.90 | 0.00E+00 | 0.00E+00 | 1.29 |
| ***CCNA_02816*** | hypothetical protein | 53.83 | 3.58 | 1.84 | 0.00E+00 | 0.00E+00 | 2.69 |
| ***CCNA_02889*** | FKBP-type peptidyl-prolyl cis-trans isomerase | 3.22 | 3.48 | 1.80 | 0.00E+00 | 0.00E+00 | 2.55 |
| ***CCNA_02820*** | TadG-family protein | 6.20 | 3.35 | 1.74 | 0.00E+00 | 0.00E+00 | 3.23 |
| ***CCNA_02879*** | genetic exchange related protein | 1.56 | 3.34 | 1.74 | 2.13E-03 | 1.69E-02 | 2.08 |
| ***CCNA_00085*** | dienelactone hydrolase-related protein | 5.55 | 3.28 | 1.71 | 0.00E+00 | 0.00E+00 | 1.01 |
| ***CCNA_03711*** | ribosome-associated factor Y | 38.90 | 3.24 | 1.70 | 0.00E+00 | 0.00E+00 | 2.42 |
| ***CCNA_00687*** | hypothetical protein | 5.58 | 3.12 | 1.64 | 0.00E+00 | 0.00E+00 | 2.20 |
| ***CCNA_02308*** | hypothetical protein | 2.04 | 2.98 | 1.58 | 0.00E+00 | 0.00E+00 | 1.74 |
| ***CCNA_02846*** | DegP/HtrA-family serine protease | 185.61 | 2.98 | 1.57 | 0.00E+00 | 0.00E+00 | 3.23 |
| ***CCNA_00734*** | GumN superfamily protein | 8.48 | 2.95 | 1.56 | 0.00E+00 | 0.00E+00 | 2.73 |
| ***rpoH*** | RNA polymerase sigma factor RpoH | 93.27 | 2.79 | 1.48 | 0.00E+00 | 0.00E+00 | 1.83 |
| ***CCNA_03601*** | hemolysin III-related membrane protein | 21.88 | 2.70 | 1.43 | 0.00E+00 | 0.00E+00 | 3.17 |
| ***CCNA_03602*** | patatin family phospholipase domain protein | 9.61 | 2.69 | 1.43 | 0.00E+00 | 0.00E+00 | 2.07 |
| ***CCNA_02721*** | peptidase, M16 family | 38.79 | 2.64 | 1.40 | 0.00E+00 | 0.00E+00 | 1.63 |
| ***CCNA_02781*** | hypothetical protein | 7.06 | 2.63 | 1.39 | 0.00E+00 | 0.00E+00 | 1.32 |
| ***CCNA_03083*** | hybrid sensor histidine kinase/receiver domain protein | 4.07 | 2.61 | 1.38 | 0.00E+00 | 0.00E+00 | 1.43 |
| ***tolB*** | Tol-Pal system protein TolB | 37.49 | 2.56 | 1.36 | 0.00E+00 | 0.00E+00 | 2.21 |

| | | | | | | | |
|---|---|---|---|---|---|---|---|
| ***CCNA_02780*** | hypothetical protein | 13.43 | 2.54 | 1.34 | 0.00E+00 | 0.00E+00 | 1.34 |
| ***CCNA_00378*** | thiol:disulfide interchange protein dsbA | 38.59 | 2.48 | 1.31 | 0.00E+00 | 0.00E+00 | 2.13 |
| ***CCNA_01955*** | zinc metalloprotease | 43.40 | 2.47 | 1.30 | 0.00E+00 | 0.00E+00 | 1.69 |
| ***CCNA_02987*** | hypothetical protein | 2.30 | 2.43 | 1.28 | 3.21E-05 | 4.06E-04 | 0.94 |
| ***mmpA*** | membrane endopeptidase MmpA | 37.89 | 2.37 | 1.24 | 0.00E+00 | 0.00E+00 | 2.39 |
| ***CCNA_00612*** | L-asparaginase | 1.17 | 2.35 | 1.23 | 1.71E-09 | 3.69E-08 | 0.95 |
| ***CCNA_01147*** | acetyltransferase | 6.43 | 2.35 | 1.23 | 0.00E+00 | 0.00E+00 | 1.12 |
| ***CCNA_03506*** | putative transcriptional regulator | 11.75 | 2.33 | 1.22 | 0.00E+00 | 0.00E+00 | 1.69 |
| ***CCNA_00124*** | hypothetical protein | 79.70 | 2.31 | 1.21 | 0.00E+00 | 0.00E+00 | 2.61 |
| ***CCNA_01569*** | Helix-turn-helix DNA-binding protein | 0.96 | 2.31 | 1.21 | 2.17E-06 | 3.31E-05 | 1.35 |
| ***CCNA_03726*** | dTDP-4-amino-4,6-dideoxygalactose transaminase, WecE family | 26.02 | 2.27 | 1.18 | 0.00E+00 | 0.00E+00 | 0.81 |
| ***CCNA_02219*** | hypothetical protein | 188.07 | 2.24 | 1.16 | 0.00E+00 | 0.00E+00 | 1.83 |
| ***CCNA_01654*** | cyclophilin-type peptidylprolyl cis-trans isomerase | 12.73 | 2.22 | 1.15 | 0.00E+00 | 0.00E+00 | 1.69 |
| ***CCNA_03710*** | nitrogen regulatory IIA protein | 9.18 | 2.18 | 1.12 | 0.00E+00 | 0.00E+00 | 1.25 |
| ***CCNA_01724*** | Tetratricopeptide repeat protein | 126.99 | 2.13 | 1.09 | **0.00E+00** | 0.00E+00 | 1.11 |
| ***CCNA_01708*** | hypothetical protein | 3.96 | 2.12 | 1.09 | 4.83E-13 | 1.33E-11 | 1.21 |
| ***CCNA_00519*** | hypothetical protein | 60.15 | 2.08 | 1.05 | 0.00E+00 | 0.00E+00 | 1.36 |
| ***CCNA_01072*** | hypothetical protein | 1.54 | 2.07 | 1.05 | 7.11E-03 | 4.66E-02 | 0.76 |
| ***nstA*** | redox cell cycle regulatory protein NstA | 33.11 | 2.06 | 1.04 | 0.00E+00 | 0.00E+00 | 2.24 |
| ***CCNA_01759*** | peptidyl-prolyl cis-trans isomerase | 148.97 | 2.05 | 1.03 | 0.00E+00 | 0.00E+00 | 2.37 |
| ***CCNA_01653*** | cyclophilin-type peptidylprolyl cis-trans isomerase | 57.29 | 2.04 | 1.03 | 0.00E+00 | 0.00E+00 | 2.17 |
| ***CCNA_01583*** | phosphate-binding protein | 20.27 | 2.04 | 1.03 | 0.00E+00 | 0.00E+00 | 2.18 |
| ***CCNA_03454*** | UDP-2,3-diacylglucosamine hydrolase | 11.65 | 2.03 | 1.02 | 0.00E+00 | 0.00E+00 | 0.85 |
| ***rsaFa*** | type I secretion outer membrane protein RsaFa | 61.53 | 2.02 | 1.02 | 0.00E+00 | 0.00E+00 | 1.75 |
| ***smc*** | chromosome partition protein Smc | 7.42 | 1.97 | 0.98 | 0.00E+00 | 0.00E+00 | 1.45 |
| ***CCNA_00614*** | short-chain fatty acids transporter | 1.81 | 1.96 | 0.97 | 2.97E-12 | 7.67E-11 | 0.59 |
| ***CCNA_02336*** | lysophospholipase L2 | 2.41 | 1.96 | 0.97 | 5.31E-10 | 1.16E-08 | 1.20 |
| ***CCNA_00290*** | autotransporter outer membrane beta-barrel domain-containing protein | 154.14 | 1.94 | 0.95 | 0.00E+00 | 0.00E+00 | 1.68 |

| | | | | | | | |
|---|---|---|---|---|---|---|---|
| ***CCNA_02106*** | TonB-dependent outer membrane receptor | 3.86 | 1.94 | 0.95 | 0.00E+00 | 0.00E+00 | 3.83 |
| ***CCNA_00224*** | TonB-dependent receptor protein | 2.55 | 1.93 | 0.95 | 2.44E-15 | 7.75E-14 | 7.30 |
| ***CCNA_00615*** | TonB-dependent receptor | 7.90 | 1.88 | 0.91 | 0.00E+00 | 0.00E+00 | 1.21 |
| ***CCNA_01497*** | ADP-L-glycero-D-manno-heptose-6-epimerase | 9.56 | 1.88 | 0.91 | 0.00E+00 | 0.00E+00 | 1.56 |
| ***CCNA_02337*** | nonspecific lipid-transfer protein | 99.02 | 1.87 | 0.91 | 0.00E+00 | 0.00E+00 | 1.45 |
| ***CCNA_02617*** | histidinol-phosphate transaminase | 12.53 | 1.87 | 0.90 | 1.59E-10 | 3.57E-09 | 0.93 |
| ***CCNA_01496*** | 2-dehydro-3-deoxyphosphooctonate aldolase | 61.42 | 1.87 | 0.90 | 0.00E+00 | 0.00E+00 | 1.57 |
| ***CCNA_03291*** | WecE-family pyridoxal phosphate-dependent enzyme | 1.28 | 1.87 | 0.90 | 1.05E-07 | 1.88E-06 | 1.15 |
| ***CCNA_00379*** | thiol:disulfide interchange protein dsbA | 89.18 | 1.87 | 0.90 | 0.00E+00 | 0.00E+00 | 1.58 |
| ***CCNA_00617*** | arginine N-succinyltransferase, beta chain | 3.36 | 1.86 | 0.90 | 1.34E-13 | 3.76E-12 | 1.44 |
| ***CCNA_00218*** | conserved hypothetical protein | 12.95 | 1.86 | 0.90 | 7.51E-12 | 1.92E-10 | 0.96 |
| ***CCNA_00486*** | TonB-dependent receptor | 1.17 | 1.86 | 0.89 | 9.21E-15 | 2.75E-13 | 1.32 |
| ***pdxA*** | 4-hydroxythreonine-4-phosphate dehydrogenase | 17.11 | 1.81 | 0.86 | 0.00E+00 | 0.00E+00 | 2.37 |
| ***CCNA_00061*** | cytochrome p450 | 3.04 | 1.80 | 0.85 | 3.49E-11 | 8.57E-10 | 1.08 |
| ***CCNA_00782*** | aminobenzoyl-glutamate transport protein | 12.27 | 1.80 | 0.85 | 0.00E+00 | 0.00E+00 | 2.88 |
| ***CCNA_00138*** | TonB-dependent receptor | 2.31 | 1.78 | 0.83 | 4.46E-08 | 8.40E-07 | 0.65 |
| ***CCNA_03461*** | hypothetical protein | 40.08 | 1.76 | 0.82 | 0.00E+00 | 0.00E+00 | 7.72 |
| ***bamB*** | beta-barrel assembly machine (BAM) protein BamB | 140.54 | 1.76 | 0.82 | 0.00E+00 | 0.00E+00 | 1.84 |
| ***sgt1*** | sphingolipid glucuronosyltransferase | 1.68 | 1.76 | 0.82 | 5.84E-09 | 1.19E-07 | 1.21 |
| ***CCNA_03773*** | DUF3597 domain-containing protein | 6.08 | 1.76 | 0.81 | 0.00E+00 | 0.00E+00 | 1.62 |
| ***CCNA_01592*** | SN-glycerol-3-phosphate transport ATP-binding protein | 74.84 | 1.75 | 0.81 | 0.00E+00 | 0.00E+00 | 2.44 |
| ***CCNA_03460*** | hypothetical protein | 47.24 | 1.71 | 0.77 | 0.00E+00 | 0.00E+00 | 4.95 |
| ***zauP*** | cell division protein ZauP | 31.35 | 1.70 | 0.76 | 0.00E+00 | 0.00E+00 | 0.75 |
| ***CCNA_03097*** | aldo/keto reductase family protein | 7.47 | 1.70 | 0.76 | 0.00E+00 | 0.00E+00 | 1.36 |
| ***CCNA_00897*** | hypothetical protein | 45.15 | 1.69 | 0.76 | 1.78E-10 | 3.96E-09 | 1.84 |
| ***CCNA_01956*** | DsbA Com1-like subfamily protein | 45.47 | 1.68 | 0.75 | 0.00E+00 | 0.00E+00 | 1.58 |
| ***CCNA_01757*** | dimethyladenosine transferase | 5.82 | 1.68 | 0.75 | 6.39E-11 | 1.52E-09 | 2.10 |
| ***bamE*** | beta-barrel assembly machine (BAM) protein BamE | 61.11 | 1.66 | 0.73 | 0.00E+00 | 0.00E+00 | 1.88 |

| | | | | | | | |
|---|---|---|---|---|---|---|---|
| ***CCNA_01020*** | LacI-family transcriptional regulator | 11.66 | 1.65 | 0.73 | 0.00E+00 | 0.00E+00 | 1.80 |
| ***CCNA_01073*** | zinc-finger protein | 8.27 | 1.65 | 0.72 | 1.74E-09 | 3.74E-08 | 1.15 |
| ***CCNA_02433*** | hypothetical protein | 18.24 | 1.65 | 0.72 | 1.20E-11 | 2.98E-10 | 3.51 |
| ***bamA*** | beta-barrel assembly machine (BAM) protein BamA | 131.43 | 1.65 | 0.72 | 0.00E+00 | 0.00E+00 | 1.51 |
| ***bamD*** | beta-barrel assembly machine (BAM) protein BamD | 135.25 | 1.64 | 0.72 | 0.00E+00 | 0.00E+00 | 1.40 |
| ***CCNA_00907*** | iron-containing redox enzyme | 6.98 | 1.64 | 0.72 | 1.45E-13 | 4.02E-12 | 1.22 |
| ***ybgF*** | Tol-Pal system periplasmic component YbgF | 14.70 | 1.64 | 0.71 | 0.00E+00 | 0.00E+00 | 1.31 |
| ***CCNA_00529*** | hypothetical protein | 3.12 | 1.63 | 0.71 | 3.63E-04 | 3.65E-03 | 1.41 |
| ***bamF*** | beta-barrel assembly machine (BAM) protein BamF | 22.26 | 1.63 | 0.70 | 6.44E-13 | 1.75E-11 | 1.29 |
| ***CCNA_00307*** | phospholipid-lipopolysaccharide ABC transporter | 14.67 | 1.63 | 0.70 | 0.00E+00 | 0.00E+00 | 1.54 |
| ***CCNA_03806*** | outer membrane lipoprotein | 84.21 | 1.61 | 0.69 | 1.43E-13 | 4.01E-12 | 2.16 |
| ***CCNA_02594*** | heat shock endopeptidase HtpX | 15.68 | 1.61 | 0.69 | 9.53E-11 | 2.21E-09 | 0.85 |
| ***CCNA_01088*** | conserved hypothetical membrane protein | 10.20 | 1.61 | 0.69 | 6.17E-13 | 1.69E-11 | 1.68 |
| ***tolQ*** | TolQ protein | 43.62 | 1.59 | 0.67 | 0.00E+00 | 0.00E+00 | 1.44 |
| ***CCNA_02830*** | DNA repair protein RadC | 4.17 | 1.58 | 0.66 | 4.89E-03 | 3.43E-02 | 0.65 |
| ***CCNA_01344*** | hypothetical protein | 5.53 | 1.58 | 0.66 | 3.33E-09 | 7.02E-08 | 1.23 |
| ***lpxC*** | UDP-3-O-(3-hydroxymyristoyl) N-acetylglucosaminedeacetylase | 10.50 | 1.58 | 0.66 | 1.65E-14 | 4.87E-13 | 0.49 |
| ***CCNA_01636*** | alkaline phosphatase | 19.82 | 1.56 | 0.64 | 0.00E+00 | 0.00E+00 | 1.84 |
| ***bacB*** | bactofilin B BacB | 20.62 | 1.56 | 0.64 | 1.13E-14 | 3.36E-13 | 0.95 |
| ***CCNA_00858*** | TonB-dependent receptor | 36.29 | 1.56 | 0.64 | 0.00E+00 | 0.00E+00 | 30.90 |
| ***CCNA_00784*** | peptidoglycan-associated outer membrane lipoprotein | 10.61 | 1.56 | 0.64 | 4.66E-15 | 1.42E-13 | 1.47 |
| ***CCNA_03838*** | gluconate 2-dehydrogenase/glyoxylate reductase/hydroxypyruvate reductase | 24.24 | 1.55 | 0.63 | 0.00E+00 | 0.00E+00 | 1.31 |
| ***CCNA_02571*** | major facilitator superfamily transporter | 0.92 | 1.54 | 0.63 | 9.72E-05 | 1.14E-03 | 1.15 |
| ***CCNA_02743*** | TonB dependent receptor | 95.42 | 1.54 | 0.62 | 0.00E+00 | 0.00E+00 | 3.43 |
| ***CCNA_03437*** | hypothetical protein | 2.17 | 1.53 | 0.62 | 2.54E-03 | 1.96E-02 | 0.89 |
| ***CCNA_03158*** | FAD:protein FMN transferase ApbE | 4.70 | 1.52 | 0.61 | 8.86E-07 | 1.41E-05 | 0.21 |
| ***CCNA_01092*** | ACT domain protein | 11.82 | 1.52 | 0.60 | 4.06E-06 | 5.93E-05 | 1.63 |
| ***CCNA_02859*** | hypothetical protein | 3.33 | 1.52 | 0.60 | 1.16E-03 | 9.98E-03 | 0.40 |

| | | | | | | | |
|---|---|---|---|---|---|---|---|
| ***CCNA_02979*** | aminotransferase class V-fold PLP-dependent enzyme | 3.29 | 1.51 | 0.60 | 2.25E-06 | 3.42E-05 | 1.06 |
| ***CCNA_03909*** | hypothetical protein | 48.54 | 1.51 | 0.59 | 0.00E+00 | 0.00E+00 | |
| ***CCNA_00967*** | TetR-family transcriptional regulator | 11.31 | -1.51 | -0.60 | 6.40E-11 | 1.52E-09 | 2.27 |
| ***ybgT*** | cyd operon protein YbgT | 19.78 | -1.52 | -0.61 | 3.03E-05 | 3.89E-04 | 1.08 |
| ***CCNA_03374*** | hypothetical protein | 6.95 | -1.54 | -0.62 | 1.03E-05 | 1.41E-04 | 0.68 |
| ***CCNA_01100*** | acylamino-acid-releasing enzyme | 5.58 | -1.55 | -0.63 | 8.94E-14 | 2.54E-12 | 0.85 |
| ***CCNA_02739*** | Rve-family integrase core domain protein | 23.72 | -1.56 | -0.65 | 0.00E+00 | 0.00E+00 | |
| ***CCNA_01258*** | conserved hypothetical membrane protein | 1.62 | -1.57 | -0.65 | 3.61E-04 | 3.64E-03 | 1.12 |
| ***CCNA_03089*** | YecT-family protein | 111.57 | -1.57 | -0.65 | 1.96E-08 | 3.90E-07 | 0.90 |
| ***CCNA_00660*** | transposase | 23.09 | -1.59 | -0.66 | 0.00E+00 | 0.00E+00 | 0.62 |
| ***CCNA_01121*** | MA superfamily peptidase | 7.41 | -1.59 | -0.67 | 3.77E-15 | 1.16E-13 | 0.86 |
| ***CCNA_02814*** | transposase | 23.09 | -1.62 | -0.70 | 0.00E+00 | 0.00E+00 | |
| ***CCNA_03168*** | hypothetical protein | 23.89 | -1.62 | -0.70 | 0.00E+00 | 0.00E+00 | 0.75 |
| ***CCNA_02677*** | glutathione S-transferase | 1.07 | -1.74 | -0.80 | 2.52E-03 | 1.95E-02 | 1.25 |
| ***CCNA_01202*** | membrane alanine aminopeptidase | 36.00 | -1.74 | -0.80 | 0.00E+00 | 0.00E+00 | 0.66 |
| ***CCNA_02741*** | Gcw/chp-family protein | 483.12 | -1.79 | -0.84 | 0.00E+00 | 0.00E+00 | 2.88 |
| ***CCNA_03166*** | ArsR-family transcriptional regulator | 0.86 | -1.79 | -0.84 | 4.44E-03 | 3.19E-02 | 0.91 |
| ***CCNA_02740*** | transposase | 6.42 | -1.80 | -0.85 | 5.74E-11 | 1.38E-09 | |
| ***CCNA_02813*** | transposase | 6.50 | -1.86 | -0.90 | 2.56E-12 | 6.69E-11 | |
| ***CCNA_02813*** | tRNA-Leu | 9.75 | -1.90 | -0.93 | 2.70E-04 | 2.77E-03 | |
| ***CCNA_R0145*** | small non-coding RNA | 2.81 | -1.90 | -0.93 | 2.34E-04 | 2.45E-03 | |
| ***CCNA_00661*** | transposase, HTH-Tnp1 superfamily | 6.97 | -2.01 | -1.01 | 1.30E-12 | 3.49E-11 | 0.91 |
| ***CCNA_00974*** | TonB-dependent outer membrane receptor | 169.22 | -3.48 | -1.80 | 0.00E+00 | 0.00E+00 | 1.24 |
| ***chvT*** | TonB-dependent outer membrane receptor ChvT | 808.92 | -3.85 | -1.95 | 0.00E+00 | 0.00E+00 | 1.52 |

### Table S2: Strains and plasmids used in this work.

| ***E. coli* strains with plasmids** | | | |
|---|---|---|---|
| **Strain #** | **Genotype** | **Comments** | **Source** |
| FC929 | TOP10 | Cloning strain | Invitrogen |
| FC3 | MT607 / pRK600 | Helper strain for tri-parental matings | (4) |
| ***Caulobacter* expression plasmids** | | | |
| MTLS4259 | TOP10 / pMT585 (pXGFPC-2) | Integrates at the xylose locus, $Kan^R$ | (5) |
| FC3460 | TOP10 / pMT585-*chvI* | Cloning Primers:<br>5'-gttgcatATGGCCGCGATCACGCTC-3'<br>5'-caacgagctctcaGGCTTCGCGATAAC-3' | This work |
| FC3461 | TOP10 / pMT585-*chvG* | Cloning Primers:<br>5'-gttgcatATGGCTACCGTTATCGCG-3'<br>5'-caacgagctctcaTTCGCGCGCGTCC-3' | This work |
| FC3464 | TOP10 / pMT585-*chvI(D52E)* | Generated from FC3446 | This work |
| FC3465 | TOP10 / pMT585-*chvG(H309A)* | Generated from FC3447 | This work |
| FC3466 | TOP10 / pMT585-*ntrX* | Cloning Primers:<br>5'-gttgcatATGAGCGCCGACGTTC-3'<br>5'-CaacgagctcctaCTCTTCCTCATC-3' | This work |
| FC3467 | TOP10 / pMT585-*ntrX(A19V)* | Generated from FC3452 | This work |
| FC3468 | TOP10 / pMT585-*ntrX(D53A)* | Generated from FC3452 | This work |
| FC3469 | TOP10 / pMT585-*ntrY* | Cloning Primers:<br>5'-gttgcatATGTCTTCAGTGGC-3'<br>5'-cattggtacctcaTATCATCTCCTC-3' | This work |
| FC3470 | TOP10 / pMT585-*ntrY(L70H)* | Generated from FC3455 | This work |
| FC3471 | TOP10 / pMT585-*ntrY(A123V)* | Generated from FC3455 | This work |
| FC3472 | TOP10 / pMT585-*ntrZ* | *CCNA_03863*, Cloning Primers:<br>5'-gttgcatATGGTCGCGACGCG-3'<br>5'-caacgagctcttaGAACTTGAAGGC-3' | This work |
| FC3473 | TOP10 / pMT585-*ntrZ(I99N)* | Generated from FC3458 | This work |
| FC3474 | TOP10 / pMT585-*ntrZ(Y92C)* | Generated from FC3458 | This work |
| FC3475 | TOP10 / pMT585-*chvT* | Cloning Primers:<br>5'-gttgcatATGGCGTTGAAAACCAAGG-3'<br>5'-gccgagatctttaCATCTTGTAGTTGAAGCC-3' | This work |
| **Allele Replacement Plasmids** | | | |
| FC55 | DH10B / pNPTS138 | Allele replacement plasmid, $Kan^R$, SacB | M.R.K. Alley |
| FC3476 | TOP10/ pNPTS138-Δ*chvI* | knockout allele contains first 4 and last 16 codons of *chvI*. 5' end of ORF (5'- ATGGCCGCGATC…), 3' end of ORF (…GAAGCCtga-3') | This work |
| FC3477 | TOP10 / pNPTS138-Δ*chvG* | knockout allele contains first 16 and last 10 codons of *chvG*. 5' end of ORF (5'-TTGGCTACC…), 3' end of ORF (…CGCGAAtga-3' | This work |
| FC3478 | TOP10 / pNPTS138-Δ*ntrX* | knockout allele contains first 12 and last 12 codons of *ntrX*. 5' end of ORF (5'-ATGAGCGCC…), 3' end of ORF (…GAAGAGtag-3') | This work |
| FC3479 | TOP10 / pNPTS138-Δ*ntrY* | knockout allele contains first 10 and last 15 codons of *ntrY*. 5' end of ORF (5'-ATGTCTTCA…), 3' end of ORF (…ATGATAtga-3') | This work |
| FC3480 | TOP10 / pNPTS138-Δ*ntrZ* | knockout allele contains first 4 and last 12 codons of *ntrZ* (*CCNA_03863)*. 5' end of ORF (5'-ATGGTCGCG…), 3' end of ORF (…AAGTTCtaa-3') | This work |
| FC2373 | TOP10 / pNPTS138-Δ*chvT* | Knockout allele contains first 2 and lasts 16 codons of *chvT* based on an outdated annotation. 5' end of ORF (5'-TTGCTC…), 3' end of ORF s(…AAGATGTAA-3') | Ola Fergin |
| FC3481 | TOP10 / pNPTS138-*ntrX* | Allele-replacement construct contains ~500 bp upstream, *ntrX*, and ~500 bp downstream | This work |
| FC3482 | TOP10 / pNPTS138-*ntrX(A19V)* | Allele-replacement construct contains ~500 bp upstream, *ntrX(A19V)*, and ~500 bp downstream | This work |
| FC3483 | TOP10 / pNPTS138-*chvI* | Allele-replacement construct contains ~500 bp upstream, *chvI*, and ~500 bp downstream | This work |
| FC3484 | TOP10 / pNPTS138-*chvI(D52A)* | Allele-replacement construct contains ~500 bp upstream, *chvI(D52A)*, and ~500 bp downstream | This work |
| FC3485 | TOP10 / pNPTS138-*chvI(D52N)* | Allele-replacement construct contains ~500 bp upstream, *chvI(D52N)*, and ~500 bp downstream | This work |
| FC3486 | TOP10 / pNPTS138-*chvI(D52E)* | Allele-replacement construct contains ~500 bp upstream, *chvI(D52E)*, and ~500 bp downstream | This work |

| | | | |
|---|---|---|---|
| FC3560 | TOP10 / pNPTS138-*ntrX-HA* | Allele-replacement construct contains last ~500 bp of *ntrX*, *HA* tag, and ~500 bp downstream of *ntrX* | This work |
| **Transcriptional *lacZ* fusion plasmids** | | | |
| FC3487 | TOP10 / pRKlac290-$P_{CCNA\_03987}$ | Plasmid containing $P_{CCNA\_03987}$-*lacZ* transcriptional fusion, tet$^r$.<br>Cloning Primers (430 bp upstream of ORF):<br>5'-ggatgaattcccttcttggtgtcgc-3'<br>5'-gaacaagcttttgtaatgtccgagtttc-3' | This work |
| FC3488 | TOP10 / pRKlac290-$P_{CCNA\_00889}$ | Plasmid containing $P_{CCNA\_00889}$-*lacZ* transcriptional fusion, tet$^r$.<br>Cloning Primers (500 bp upstream of ORF):<br>5'-ggatgaattccgtcggaccgctatg-3'<br>5'-gaacaagcttacgcgtctccaaggg-3' | This work |
| FC3489 | TOP10 / pRKlac290-$P_{CCNA\_01341}$ | Plasmid containing $P_{CCNA\_01341}$-*lacZ* transcriptional fusion, tet$^r$.<br>Cloning Primers (500 bp upstream of ORF):<br>5'-ggatgaattcgtgaagcggttgggtg-3'<br>5'-gaacaagcttgaaggactcctagc-3' | This work |
| FC3490 | TOP10 / pRKlac290-$P_{chvR}$ | Plasmid containing $P_{chvR}$-*lacZ* transcriptional fusion, tet$^r$. | (6) |
| FC3491 | TOP10 / pRKlac290-$P_{nstA}$ | Plasmid containing $P_{nstA}$-*lacZ* transcriptional fusion, tet$^r$.<br>Cloning Primers (500 bp upstream of ORF):<br>5'-ggatgaattccgcaggtgcggccgc-3'<br>5'-gaacaagcttgggaactgtccgctc-3' | This work |
| FC3492 | TOP10 / pRKlac290-$P_{chvI}$ | Plasmid containing $P_{chvI}$-*lacZ* transcriptional fusion, tet$^r$.<br>Cloning Primers (500 bp upstream of ORF):<br>5'-ggatgaattcgtgtagagatcggcg-3'<br>5'-gaactctagacgggtcgatacgtgtc-3' | This work |
| ***Caulobacter crescentus* strains** | | | |
| **Strain #** | **Genotype** | **Source** | **Figure** |
| FC19 | Wild type CB15 | (7) | 2, 3, 4, 5, S3, S4 |
| FC3493 | WT CB15 (Allele replaced from Δ*chvI*) | This work | 2 |
| FC648 | CB15 *xylX*::pMT585 | This work | 1, 2, 4, 5, 7, S1, S3, S6 |
| FC3494 | CB15 *xylX*::pMT585-*ntrX* | This work | 4 |
| FC3495 | CB15 *xylX*::pMT585-*ntrX(D53A)* | This work | 4 |
| FC3496 | CB15 *chvI(D52A)* | This work | 2, S3 |
| FC3497 | CB15 *chvI(D52N)* | This work | 2, S3 |
| FC3498 | CB15 *chvI(D52E)* | This work | 2, S3 |
| FC3499 | CB15 Δ*chvI* | This work | 2, 3, 4, 5, S2, S3, S4 |
| FC3500 | CB15 Δ*chvI* (Allele replaced from Δ*chvI* Δ*ntrX)* | This work | 4 |
| FC3501 | CB15 Δ*chvI ntrX(A19V)* | This work | 4 |
| FC3502 | CB15 Δ*chvI xylX*::pMT585 | This work | 1, 2, 4, 5, 7, S1, S3, S6 |
| FC3503 | CB15 Δ*chvI xylX*::pMT585-*chvI* | This work | 1, S1 |
| FC3504 | CB15 Δ*chvI xylX*::pMT585-*chvI(D52E)* | This work | 2, S3 |
| FC3505 | CB15 Δ*chvI xylX*::pMT585-*ntrX* | This work | 4 |
| FC3506 | CB15 Δ*chvI xylX*::pMT585-*ntrX(A19V)* | This work | 4 |
| FC3507 | CB15 Δ*chvI xylX*::pMT585-*ntrX(D53A)* | This work | 4 |
| FC3508 | CB15 Δ*chvI xylX*::pMT585-*ntrY* | This work | 5 |
| FC3509 | CB15 Δ*chvI xylX*::pMT585-*ntrY(L70H)* | This work | 5 |
| FC3510 | CB15 Δ*chvI xylX*::pMT585-*ntrY(A123V)* | This work | 5 |
| FC3511 | CB15 Δ*chvI xylX*::pMT585-*ntrZ* | This work | 5 |
| FC3512 | CB15 Δ*chvI xylX*::pMT585-*ntrZ(I99N)* | This work | 5 |
| FC3513 | CB15 Δ*chvI xylX*::pMT585-*ntrZ(Y92C)* | This work | 5 |
| FC3514 | CB15 Δ*chvG* | This work | |
| FC3515 | CB15 Δ*chvG xylX*::pMT585 | This work | 1, 2, S1, S3 |
| FC3516 | CB15 Δ*chvG xylX*::pMT585-*chvG* | This work | 1, S1 |
| FC3517 | CB15 Δ*chvG xylX*::pMT585-*chvG(H309A)* | This work | 2 |
| FC3518 | CB15 Δ*chvG xylX*::pMT585-*chvI(D52E)* | This work | 2, S3 |
| FC3519 | CB15 Δ*ntrX* | This work | 4 |
| FC3520 | CB15 Δ*ntrY* | This work | 5 |
| FC3521 | CB15 Δ*ntrY xylX*::pMT585 | This work | 5 |
| FC3522 | CB15 Δ*ntrY xylX*::pMT585-*ntrY* | This work | 5 |
| FC3523 | CB15 Δ*ntrY xylX*::pMT585-*ntrZ* | This work | 5 |
| FC3524 | CB15 Δ*ntrY xylX*::pMT585-*ntrZ(I99N)* | This work | 5 |
| FC3526 | CB15 Δ*ntrZ* | This work | 5 |
| FC3527 | CB15 Δ*ntrZ xylX*::pMT585 | This work | 5 |
| FC3528 | CB15 Δ*ntrZ xylX*::pMT585-*ntrZ* | This work | 5 |

| | | | |
|---|---|---|---|
| FC3529 | CB15 Δ*ntrZ xylX*::pMT585-*ntrY* | This work | 5 |
| FC3530 | CB15 Δ*ntrZ xylX*::pMT585-*ntrY(L70H)* | This work | 5 |
| FC3531 | CB15 Δ*ntrY* Δ*ntrZ* | This work | 5 |
| FC3532 | CB15 Δ*chvI* Δ*ntrX* | This work | 4 |
| FC3533 | CB15 Δ*chvT xylX*::pMT585 | This work | 7, S6 |
| FC3534 | CB15 Δ*chvI* Δ*chvT* | This work | |
| FC3535 | CB15 Δ*chvI* Δ*chvT xylX*::pMT585 | This work | 7, S6 |
| FC3536 | CB15 Δ*chvI* Δ*chvT xylX*::pMT585-*chvT* | This work | 7, S6 |
| FC3537 | CB15 / pRKlac290-$P_{CCNA\_03987}$ | This work | 6 |
| FC3538 | CB15 / pRKlac290-$P_{CCNA\_00889}$ | This work | 6 |
| FC3539 | CB15 / pRKlac290-$P_{CCNA\_01341}$ | This work | 6 |
| FC3540 | CB15 / pRKlac290-$P_{chvR}$ | This work | 6 |
| FC3541 | CB15 / pRKlac290-$P_{nstA}$ | This work | 6 |
| FC3542 | CB15 / pRKlac290-$P_{chvI}$ | This work | 6 |
| FC3543 | CB15 Δ*chvI* / pRKlac290-$P_{CCNA\_03987}$ | This work | 6 |
| FC3544 | CB15 Δ*chvI* / pRKlac290-$P_{CCNA\_00889}$ | This work | 6 |
| FC3545 | CB15 Δ*chvI* / pRKlac290-$P_{CCNA\_01341}$ | This work | 6 |
| FC3546 | CB15 Δ*chvI* / pRKlac290-$P_{chvR}$ | This work | 6 |
| FC3547 | CB15 Δ*chvI* / pRKlac290-$P_{nstA}$ | This work | 6 |
| FC3548 | CB15 Δ*chvI* / pRKlac290-$P_{chvI}$ | This work | 6 |
| FC3549 | CB15 Δ*ntrX* / pRKlac290-$P_{CCNA\_03987}$ | This work | 6 |
| FC3550 | CB15 Δ*ntrX* / pRKlac290-$P_{CCNA\_00889}$ | This work | 6 |
| FC3551 | CB15 Δ*ntrX* / pRKlac290-$P_{CCNA\_01341}$ | This work | 6 |
| FC3552 | CB15 Δ*ntrX* / pRKlac290-$P_{chvR}$ | This work | 6 |
| FC3553 | CB15 Δ*ntrX* / pRKlac290-$P_{nstA}$ | This work | 6 |
| FC3554 | CB15 Δ*ntrX* / pRKlac290-$P_{chvI}$ | This work | 6 |
| FC3561 | CB15 *ntrX-HA* | This work | 5 |
| FC3562 | CB15 Δ*ntrY ntrX-HA* | This work | 5 |
| FC3563 | CB15 Δ*ntrZ ntrX-HA* | This work | 5 |
| FC3564 | CB15 Δ*ntrY* Δ*ntrZ ntrX-HA* | This work | 5 |
